# *Blautia wexlerae* Transforms Dietary Fatty Acids to Activate Enteroendocrine Signaling and Improve Metabolic Health in Mice and Humans

**DOI:** 10.64898/2026.03.13.709143

**Authors:** Yanjia Jason Zhang, Marian Tanofsky-Kraff, Mildred M. Reyes, Daniel Zeve, Kigumbi J. Ehrmann, Jennifer Lee, Ana Paula Schaan, Ana Prado, Xiao Corey Ma, Megan N. Parker, Sheila M. Brady, Elisa Saint-Denis, Karina Sharma, Bianca Frintu, Camila Richmond, Nirav Desai, Vladimir Yeliseyev, Lynn Bry, Julian Avila-Pacheco, Clary B. Clish, Matthew Quealy, Jon Clardy, David T. Breault, Yufang Ding, Xingwen Wang, Marco Jost, Mathilde Poyet, Mathieu Groussin, Jack A. Yanovski, Wayne I. Lencer, Eric J. Alm

**Affiliations:** Center for Microbiome Informatics & Therapeutics, Massachusetts Institute of Technology, Cambridge, MA; Division of Gastroenterology, Hepatology & Nutrition, Department of Pediatrics, Boston Children’s Hospital, Boston, MA; Harvard Digestive Disease Center and Brigham and Women’s Hospital, Boston, MA; Department of Medical and Clinical Psychology, Uniformed Services University of the Health Sciences (USUHS), Bethesda, MD, 20814, USA; Section on Growth and Obesity (SGO), Division of Intramural Research, Eunice Kennedy Shriver National Institute of Child Health and Human Development (NICHD), National Institutes of Health (NIH), DHHS, Bethesda, MD 20892, USA; Division of Endocrinology, Department of Pediatrics, Boston Children’s Hospital, Boston, MA; The Jean Mayer United States Department of Agriculture Human Nutrition Research Center on Aging at Tufts University, Boston, MA; The Graduate School of Biomedical Sciences, Tufts University, Boston, MA; Tufts University School of Medicine, Boston, MA; Institute of Experimental Medicine, Kiel University, Kiel, Germany; Institute of Clinical Molecular Biology, Kiel University, Kiel, Germany; Department of Biological Chemistry and Molecular Pharmacology, Harvard Medical School and Blavatnik Institute, Boston, MA, USA; Massachusetts Host-Microbiome Center, Department of Pathology, Brigham and Women’s Hospital, Harvard Medical School, Boston, MA, USA; Metabolomics Platform, The Broad Institute of MIT and Harvard, Cambridge, MA, USA; Department of Microbiology, Harvard Medical School, Boston, MA, USA; Global Microbiome Conservancy; Amgen, 1 Amgen Center Dr, Thousand Oaks, CA 91320

## Abstract

Metabolites produced by the gut microbiome influence host metabolic health, but how this occurs remains incompletely defined. Here, we report that a common human gut commensal, *Blautia wexlerae*, converts dietary fats into bioactive metabolites that induce gut hormone production to affect glucose metabolism and suppress appetite. We found that colonization with *Blautia wexlerae* correlated with healthier eating behaviors in humans. *Blautia wexlerae* encodes a unique acyl transferase and is capable of producing acyl amines from nutrient substrates. These *Blautia* acyl amines stimulated human enteroendocrine cells to secrete GLP-1 and other gut peptide hormones more potently than endogenously produced acyl amines. When fed to mice, acyl amines improved glycemic control and decreased appetite. In humans, higher stool levels of *Blautia* DNA encoding acyl amine synthesis genes correlated with leanness and decreased dietary fat intake. These results define a mechanism of action for how *Blautia wexlerae* affects host metabolic control.

## INTRODUCTION

The gut microbiome produces a complex array of bioactive compounds that influence eating behavior, a dominant factor in the pathogenesis of obesity and its associated metabolic diseases^1–3^. In the mouse, transfer of select complex microbial communities or single microbial organisms can influence appetite and decrease the intake of palatable foods^4,5^. The mechanisms appear to involve gut-brain cross-talk, but the underlying molecular pathways linking microbial products to appetite signaling in the brain remain largely undefined^6,7^.

Gut-brain signaling via peptides secreted by enteroendocrine cells of the gut epithelium has emerged as an important lever by which gut microbes might affect both neural signaling and metabolic health^8–10^. The gut microbiome is uniquely positioned to interface with enteroendocrine cells (EEC) populating the distal gut and thus to exert influence on appetite regulation and metabolic homeostasis^11,12^. EECs secrete peptide hormones that act locally and centrally to modulate feeding behavior. Recent work indicates that coordinated activation of multiple enteroendocrine cell populations can robustly engage satiety pathways^12–19^. EECs in the distal gut, the area of highest commensal colonization, also operate critically in host glycemic control^14,20^. These observations highlight how microbes can shape gut-brain communication through the enteroendocrine system.

Here, we report that high-level colonization with the gut commensal *Blautia wexlerae* associates with healthier eating behaviors in humans. We uncover the bacterial metabolites produced by *B. wexlerae* that exert these beneficial effects by activating EECs and modulating host appetite and glucose homeostasis, establishing an integrated mechanism of action.

## RESULTS

### *Blautia wexlerae* is depleted in stools obtained from children with Loss of Control Eating

We recruited 201 healthy pediatric volunteers from an observational study (NCT02390765; Figure 1A). All participants completed an interview-based assessment for loss of control (LOC) eating, a form of defective appetite control associated with obesity, and provided stool samples for shotgun metagenomic analysis. The gut microbiomes of children with LOC eating were found to differ significantly from those without LOC eating (Figure 1B). Since only a small proportion of participants (17 individuals, 8.5%) had LOC eating, we applied the Synthetic Minority Oversampling Technique (SMOTE) to compare the gut microbiomes of participants with and without LOC eating^21^. Using species-level relative abundances, we trained a random forest classifier (accuracy: 1.0, area under the receiver operating characteristic curve (ROC): 0.98) that effectively distinguished the two groups (Figure 1B). A similar random forest classifier, developed with oversampled data for LOC eating participants, also successfully differentiated the groups (accuracy: 0.983, area under the ROC curve: 0.975, Supplemental Figure 1).

**Figure 1.**
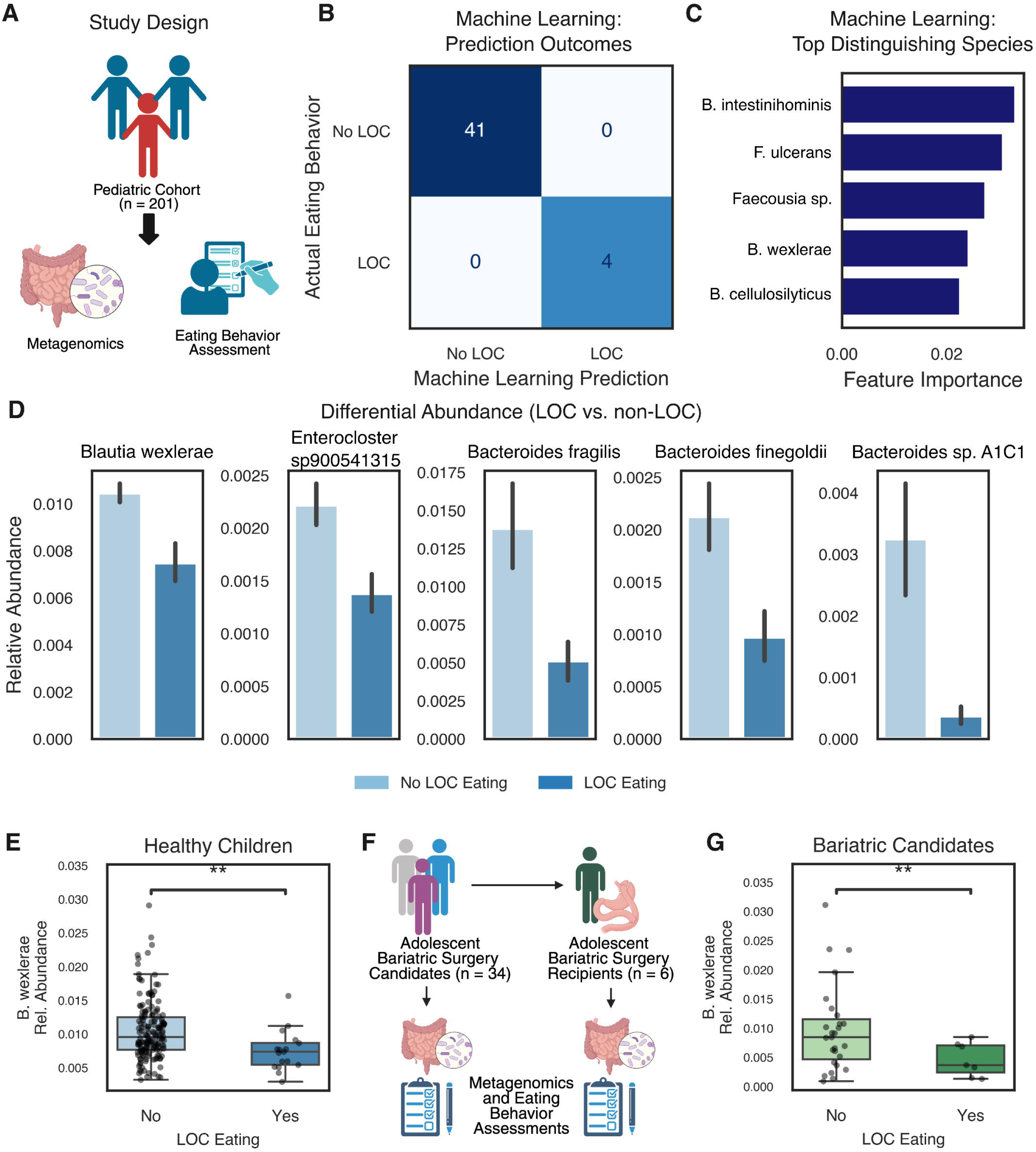
The microbiome signature of Loss of Control Eating. **A.** Schematic of the study design. 201 healthy children underwent eating behavior assessments and submitted stool for microbiome analysis using shotgun metagenomic sequencing. **B.** A confusion matrix from a machine learning algorithm using only microbiome data (relative abundance at the species level) showing accurate prediction of Loss of Control eating in children. **C.** The top five ranked features contributing to the random forest classifier, ordered by importance. **D.** The top five differentially abundant species between children with and without Loss of Control eating (unequal variance t-test, Bonferroni correction, FDR < 0.05). **E.** Healthy children with Loss of Control eating exhibited a relative depletion of *Blautia wexlerae* (**, *p* < 0.01). **F.** Schematic of the study design for adolescents seeking bariatric surgery. Thirty-four adolescents seeking bariatric surgery underwent eating behavior assessments and submitted stool for microbiome analysis using shotgun metagenomic sequencing. Six of these patients underwent bariatric surgery and submitted post-operative eating behavior assessments and microbiome samples. **G.** Adolescents seeking bariatric surgery with Loss of Control eating exhibited a relative depletion of *Blautia wexlerae* (**, *p* < 0.01).

The top microbial species distinguishing the two groups in the SMOTE-based classifier were *Barnesiella intestinihominis*, *Fusobacterium ulcerans*, *Faecousia sp., Blautia wexlerae,* and *Bacteroides cellulosilyticus* (Figure 1C). The top distinguishing species in the oversampling-based classifier were *B. wexlerae, Entercloster aldenensis, Butyricicoccus sp., Slackia_A isoflavoniconvertens,* and *Faecalibaterium sp. 4* (Supplemental Figure 1). Further analysis revealed that *B. wexlerae*, *Enterocloster sp900541315*, and three *Bacteroides* species (*B. fragilis, B. finegoldii,* and *B. sp. A1C1*) were differentially abundant between participants with and without LOC eating (unequal variance t-test, Bonferroni correction, FDR < 0.05) (Figure 1D, 1E, Supplemental Figure 2, Supplemental Table 1). Notably, all five species were less abundant in participants with LOC eating. Together, these findings demonstrate that the gut microbiome associated with LOC eating is distinct, with the depletion of *B. wexlerae* emerging as a key feature.

To test if these findings in healthy children extended to individuals with severe obesity, we recruited a smaller cohort of 34 adolescents who were seeking bariatric surgery (Figure 1F). In this case, we assessed LOC eating behavior using the LOC subscale score from the Yale Food Addiction Scale (YFAS)^22,23^. Our analysis revealed that *B. wexlerae* was also depleted in the stools of participants with LOC eating in this cohort (Figure 1G and Supplemental Figure 3).

Follow-up for this study, however, was cut short by the COVID-19 pandemic; only six participants in this prospective cohort underwent bariatric surgery, thus underpowering the study to systematically examine pre- and post-operative microbiome differences.

Still, we observed an upward trend in *B. wexlerae* relative abundance in the stools of four participants whose food addiction behaviors improved post-surgery, as measured by the total YFAS score (Supplemental Figure 4). Additionally, the two participants who did not experience an improvement in their total YFAS score exhibited a decreasing trend in *B. wexlerae* relative abundance (Supplemental Figure 4).

### *Blautia* and other related Lachnospiraceae synthesize Acyl Amines

We next aimed to uncover the mechanisms by which *Blautia* might influence host eating behaviors. Using paired fecal metabolomics and metagenomics data from the TwinsUK study, one of the largest publicly available datasets of this type (Figure 2A)^24^, we found that colonization with the genus *Blautia* was strongly associated with enrichment in metabolites from the endocannabinoid pathway (Figure 2B). Endocannabinoids include acyl amines produced endogenously by the host that are known to influence appetite^25–27^. As such, we investigated which commensal bacteria were positively or negatively associated with the fecal abundance of three key acyl amines in this metabolomic data set: oleoyl ethanolamine (OEA), palmitoyl ethanolamine (PEA), and linoleoyl ethanolamine (LEA). For all three acyl amines, the majority of operational taxonomic units (OTUs) positively associated with their abundance were classified as either *Blautia* or *Lachnospiraceae*, the taxonomic family to which the genus *Blautia* belongs (Figure 2C).

**Figure 2.**
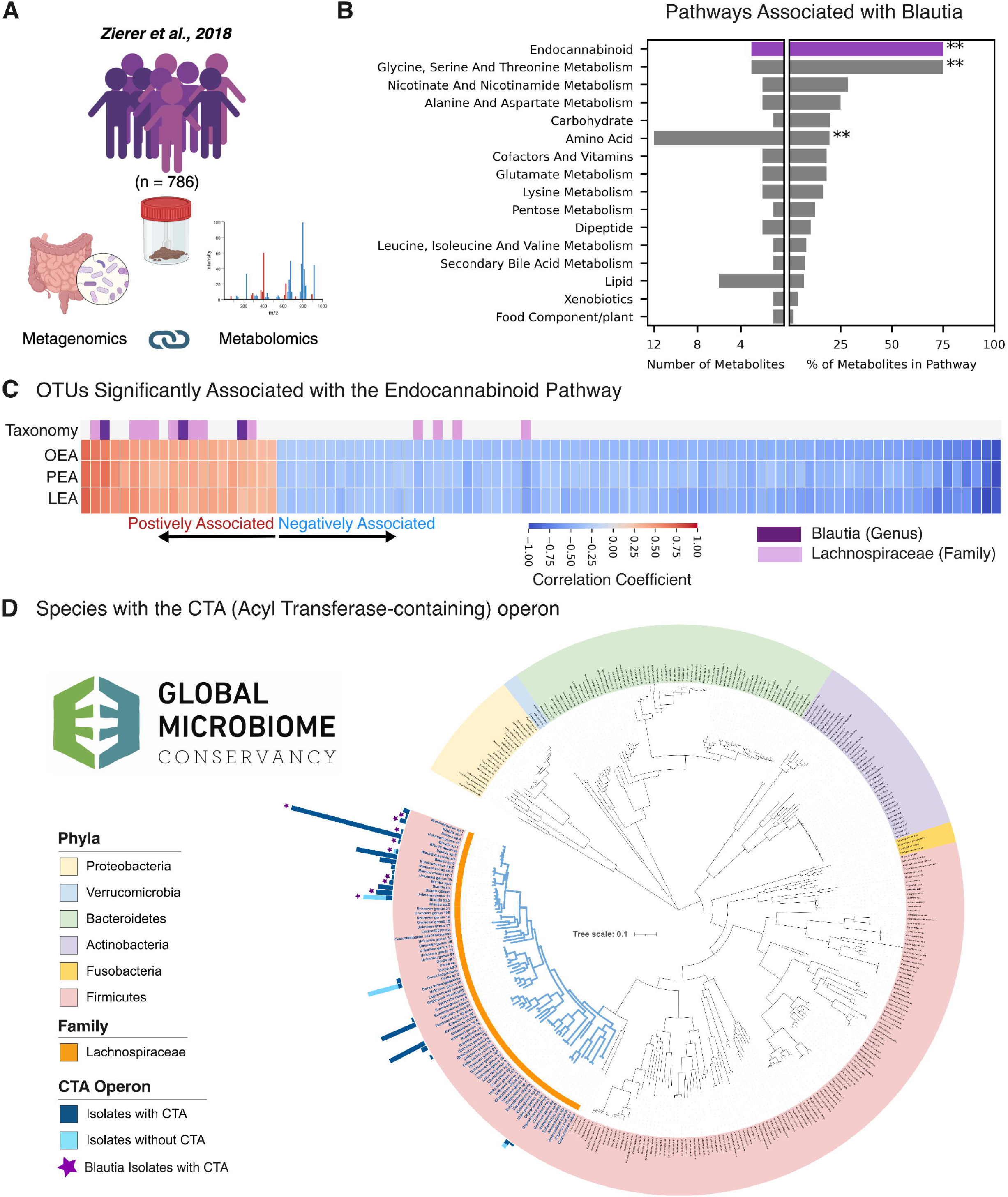
*Blautia* species synthesize acyl amines with structural homology to endogenous endocannabinoids. **A.** We utilized data and analyses from the TwinsUK cohort, which contained metagenomic and metabolomic correlations from 786 individuals. **B.** Pathways with at least one metabolite positively associated with Blautia abundance, with significantly enriched pathways marked (**, FDR < 0.01). The top pathway (purple) represented was the endocannabinoid pathway. **C.** Individual microbes (Operational Taxonomic Units, OTU) positively (red) and negatively (blue) associated with Oleoylethanolamine (OEA), Palmitoylethanolamine (PEA), and Linoleoyl Ethanolamine (LEA), three endogenously produced endocannabinoids abundance in the stool. The majority of OTUs positively associated with OEA, PEA and LEA were either *Blautia* (Purple in the Taxonomy row) or its Family *Lachnospiraceae* (Lavender in the Taxonomy row). **D.** Species in the GMbC encoding the acyl amine CTA gene operon. Dark blue bars indicate the number of isolates encoding the CTA operon, while light blue bars indicate the number of isolates without the operon (in species where at least one isolate encodes the operon). All CTA-encoding isolates are *Lachnospiraceae* (highlighted by the orange arc)*. Blautia* species encoding the CTA operon are marked by purple stars.

The strong association between *Blautia* and acyl amines observed in our TwinsUK analysis prompted us to question whether *Blautia* species might directly synthesize these metabolites. We searched the 7,781 bacterial isolate whole genomes from the Broad Institute-OpenBiome Microbiome Library (BIO-ML) and the Global Microbiome Conservancy (GMBC) biobank^28,29^ for the CTA gene cluster, a three-gene operon in *Clostridiales* species found recently to encode enzymes capable of conjugating amines with fatty acids ex vivo to produce acyl amines^30^. We found CTA operon homologs in 36 species, all of which were members of the *Lachnospiraceae* family (Figure 2D, Supplemental Table 2). Notably, all isolates of *Blautia wexlerae*—one of the most common *Lachnospiraceae* species in the human gut—encoded the entire operon (Figure 2D and Supplemental Table 2). We found both species- and strain-level variations for CTA operon encoding. For example, some *Blautia* species did not encode the operon (Figure 2D and Supplemental Table 2). At the strain level, we also found a *Coprococcus* species (another prevalent human *Lachnospiraceae*) in which one isolate encoded the operon while another did not (Figure 2D and Supplemental Table 2).

To investigate whether the CTA operon enabled acyl amine synthesis in live bacteria, we cultured four *Blautia* species from the GMBC library anaerobically in liquid media and analyzed the pelleted cells using liquid chromatography-mass spectrometry (LC-MS) (Figure 3A). Two species that encoded the CTA operon (CTA+), *B. wexlerae* and *Blautia_A sp003471165*, were compared with two species that did not (CTA-), *Blautia massiliensis* and *Blautia_A schinkii.* The *Blautia* species containing the CTA+ operon produced significantly higher levels of OEA, PEA, and LEA compared to the *Blautia* species that did not (CTA-, Figure 3B). We further tested two isolates of *Coprococcus catus*, one containing the CTA+ operon and one that did not (CTA-). We found that the CTA+ isolate had higher levels of OEA, PEA, and LEA (Supplemental Figure 5), strengthening the evidence that *Lachnospiraceae* containing the CTA operon can produce acyl amines. Notably, all the species tested encode other acyl transferases, which may account for the acyl amine synthesis observed at lower amounts in the absence of the CTA operon.

**Figure 3.**
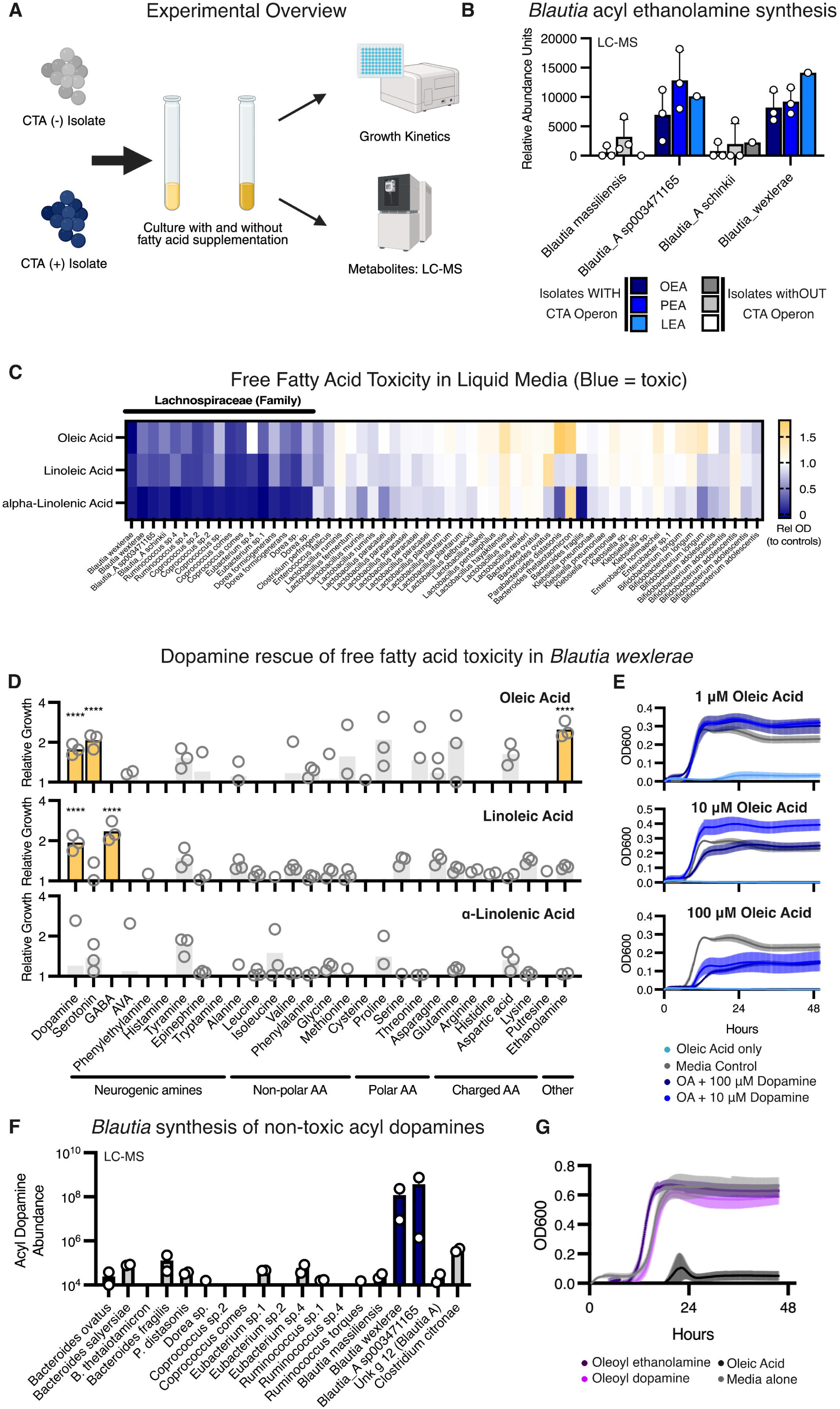
Long-chain fatty acids are toxic to *Blautia* and select amines can rescue this toxicity. **A.** Experimental scheme for growth and LC-MS measurement of acyl amines from bacterial cultures. **B.** CTA-encoding *Blautia* isolates (blue) synthesize higher amounts of OEA, PEA and LEA, measured by LC-MS, compared to *Blautia* isolates without the CTA cassette (grey). **C.** Relative OD600 of gut bacterial isolates grown with vs. without supplementation with 100 μΜ of oleic acid, linoleic acid or α-linolenic acid. Free fatty acids were common toxic to *Lachnospiraceae* isolates (black bar above the heatmap). **D.** Relative increased growth of *Blautia wexlerae* in fatty-acid enriched broth when spiked with 10 or 100 μΜ of individual amines. Relative growth was calculated as the ratio of optical densities in liquid cultures with vs. without amine supplementation, and amines that rescued growth are highlighted in yellow (****, *p* < 0.0001). **E.** Growth curves of *Blautia wexlerae* in fatty-acid enriched liquid media (top: 1 μΜ oleic acid, middle: 10 μΜ oleic acid, bottom: 100 μΜ oleic acid) with and without dopamine supplementation at 10 and 100 μΜ. Supplementation with dopamine rescued the growth of *Blautia wexlerae* in fatty-acid containing media. **F.** LC-MS measurement of acyl dopamines in bacterial cultures supplemented with oleic acid and dopamine. The two *Blautia* species encoding the CTA operon synthesized large amounts of acyl dopamines. **G.** Growth curves of *Blautia wexlerae* in liquid media supplemented with oleic acid, OEA and oleoyl dopamine. Oleic acid was toxic, while the oleoyl amines were not.

To assess the impact of exogenously applied fatty acids as substrate to enhance these reactions, we supplemented the liquid media with 100 µM free oleic acid. The *Coprococcus catus* CTA+ isolate showed a marked increase in the synthesis of several oleoyl amines, while the *Coprococcus catus* CTA- isolate showed a much smaller response (Supplemental Figure 6). Specifically, oleic acid supplementation resulted in a 4-fold increase in OEA synthesis in the CTA+ isolate, leading to a 13-fold higher OEA level compared to the CTA- isolate (Supplemental Figure 6).

### *Blautia* acyl amine synthesis links to detoxification of fatty acids

We also trialed free fatty acid supplementation in live *Blautia* species. However, when we supplemented liquid growth media with free fatty acids, we observed significant growth suppression in *Blautia* species. Supplementation with oleic acid, linoleic acid, and α-linolenic acid generally inhibited the growth of most *Blautia* species in liquid culture (Figure 3C). This inhibition was consistently found across most of the *Lachnospiraceae* species, except for a few *Coprococcus* species (Figure 3C). In contrast, we rarely observed free fatty acid toxicity in other gut-inhabiting genera, including *Lactobacillus*, *Bacteroides*, *Klebsiella*, *Enterobacteria*, and *Bifidobacterium* (Figure 3C).

These findings led us to hypothesize that *Blautia* might use acyl amine production to detoxify free fatty acids as an adaptive survival mechanism. To test this effect, we cultured *B. wexlerae* in liquid media with and without oleic acid, linoleic acid, and α-linolenic acid in a 96-deep-well format. We then screened for free amines commonly found in the human gastrointestinal tract at high concentrations (>100 µM) to determine if any of these amines provided substrate for acyl amine production. We assessed this by cultivating *B. wexlerae* in media containing both amines and toxic fatty acids, and as hypothesized, we found amines that rescued microbial growth. In growth media containing either oleic acid or linoleic acid, the addition of dopamine effectively rescued the growth of *B. wexlerae* (Figure 3D). Serotonin and ethanolamine also rescued growth in media containing oleic acid, while gamma-aminobutyric acid (GABA) rescued growth in media containing linoleic acid (Figure 3D).

To confirm these findings, we generated growth curves of *B. wexlerae* in the presence of varying concentrations of oleic acid and dopamine. Dopamine supplementation indeed restored growth in both low and high oleic acid conditions (Figure 3E). Next, we performed LC-MS on *Blautia* grown with dopamine and oleic acid, and found that, in addition to OEA, LEA, and PEA, *B. wexlerae* and *Blautia_A sp003471165* synthesized oleoyl dopamine when grown in media supplemented with oleic acid and dopamine (Figure 3F). Finally, high acyl amine concentrations (100 µM; the same level at which free fatty acids inhibited growth) in liquid media were found not toxic to *B. wexlerae* (Figure 3G). Taken together, these results provide evidence that acyl amine synthesis by *Blautia* acts to enhance survival in high-fat environments.

### *Blautia* acyl amines stimulate GLP-1 secretion from gut enteroendocrine cells and improve glycemic control

Given *B. wexlerae’s* depletion in stools from children with LOC eating, we next sought to investigate whether *Blautia*-derived acyl amines might contribute to this effect by influencing satiety signaling via the gut. Endogenously produced N-acyl ethanolamines, such as OEA, are known to induce satiety by activating enteroendocrine cells (EECs) to secrete gut peptide hormones like GLP-1^31^. We hypothesized that acyl amines exogenously produced by *Blautia* might engage similar pathways. Since GPR119 agonism by endogenous acyl amines activates EECs, we began by testing whether acyl dopamines produced by *Blautia* activated GPR119. Using a GPR119 reporter cell line, we found that oleoyl dopamine, linoleoyl dopamine, and α-linolenoyl dopamine were strong agonists of GPR119 with EC50 values in the low micromolar range and comparably more potent than the endogenously produced OEA (Figure 4A and Supplemental Figure 7). Acyl dopamines with shorter acyl chains were less potent or showed no measurable activity (Figure 4A and Supplemental Figure 7). We also tested a panel of acyl amines previously hypothesized to be bacterially derived^30^. Acyl GABA compounds exhibited no measurable GPR119 activity, while oleoyl- and linoleoyl-aminovaleric acid were modestly effective with EC50 values in the low micromolar range (Figure 4A and Supplemental Figure 7). We next stimulated the immortalized EEC cell line NCI-h716 with *Blautia* acyl amines and found that oleoyl-, linoleoyl-, and α-linolenoyl-dopamine were potent inducers of GLP-1 secretion (Figure 4B). These bacterially produced acyl dopamines induced up to three times higher GLP-1 secretion than OEA, a known GLP-1-inducing acyl amine endogenously produced (Figure 4C).

**Figure 4.**
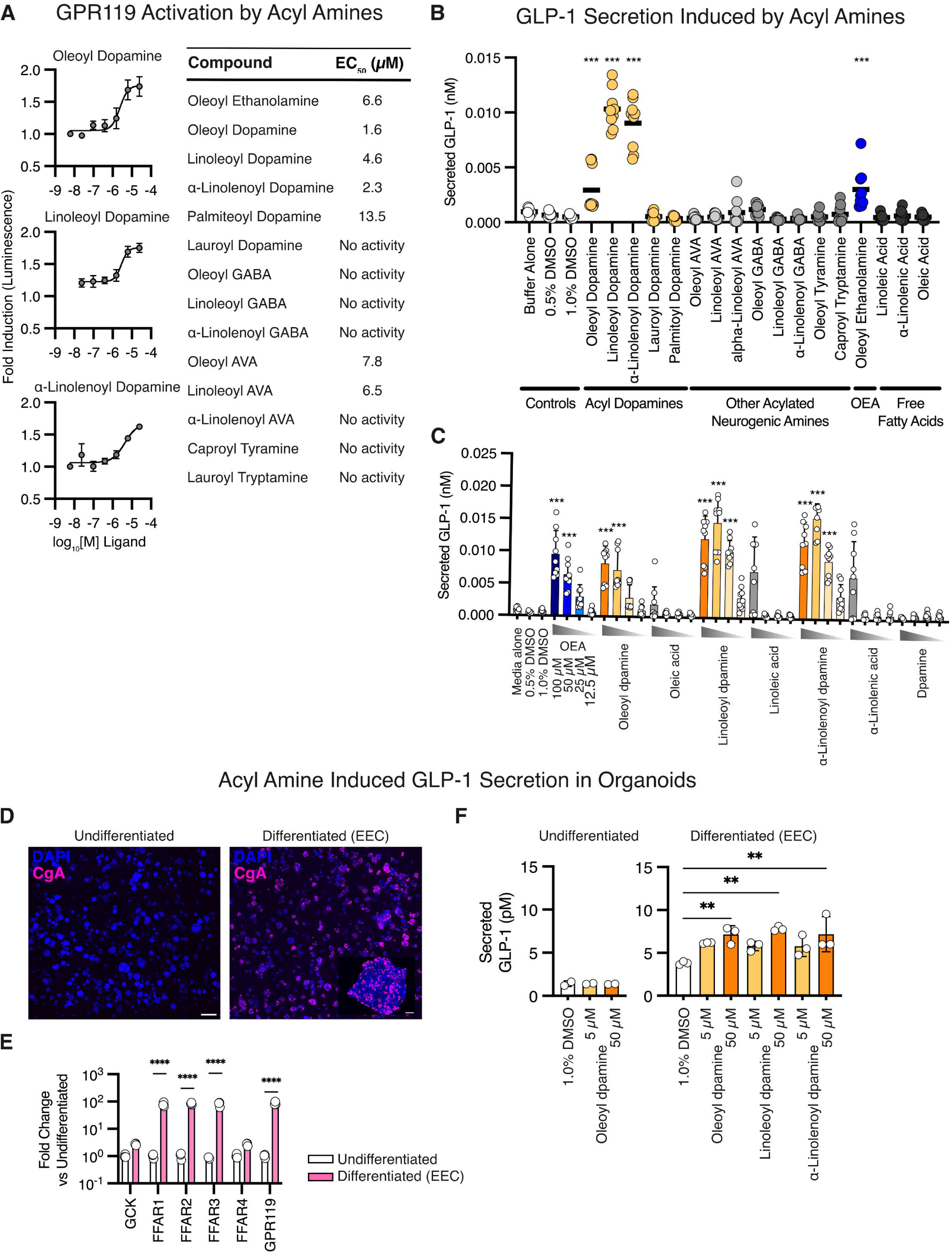
Blautia acyl amines stimulate GPR119, stimulate GLP-1 secretion and improve glycemic control in mice. **A.** Activation of a GPR119 bioreporter cell line with *Blautia* acyl amines (Left). Table of GPR119 activation EC50 values for acyl amines found in human stool. Acyl dopamines potently activated the GPR119 reporters. **B.** Secreted GLP-1 from an enteroendocrine cell line (NCI-h716) stimulated with acyl amines at 25 μΜ (***, *p* < 0.001). Acyl dopamines stimulated equal or higher amount of GLP-1 secretion compared to the known GLP-1 secretagogue OEA. **C.** Dose response for GLP-1 secretion by an enteroendocrine cells (NCI-h716) stimulated with acyl amines and controls (***, *p* < 0.001). Acyl dopamines induced more secretion of GLP-1 than the fatty acids or dopamine alone. GLP-1 secretion was measured using a luciferase-based reporter system. **D.** Organoids derived from human rectal biopsies were differentiated to include enteroendocrine cells (right) compared to controls (left), stained for EEC marker chromogranin A (CgA, magenta). Scale bar (left) = 1 mm; scale bar (right, inset) = 50 μm. **E.** Quantitative RT-PCR for free fatty acid receptors in EEC-containing vs. non-EEC containing organoids (****, *p* < 0.0001). Differentiated EEC-containing organoids expressed higher levels of free fatty acid GPCRs. **F.** Secreted GLP-1 from undifferentiated organoids (Left) and differentiated EEC-containing organoids stimulated with acyl dopamines (**, *p* < 0.01). All three acyl dopamines stimulated GLP-1 from EEC-containing organoids at 50 μΜ.

Notably, the free fatty acid precursors (oleic, linoleic, and α-linolenic acids) and dopamine alone were much less effective, or ineffective, at inducing GLP-1 secretion when applied to NCI-h716 EEC cells (Figure 4B and 4C).

To further model acyl amine stimulation of EECs, we turned to a previously developed human intestinal organoid system that amplified the differentiation of EECs in each organoid (Figure 4D)^32^. Following such EEC organoid differentiation, we observed a significant increase in the expression of fatty acid G-protein coupled receptors, including a 4-fold increase in GPR119 expression, as well as similar increases in the free fatty acid receptors FFAR1, FFAR2, and FFAR3 (Figure 4E). When we stimulated the differentiated EEC-containing intestinal organoids for two hours with oleoyl-, linoleoyl-, or α-linolenoyl-dopamine, GLP-1 secretion increased two- to three-fold (Figure 4F). Longer incubation times with bacterial acyl dopamines also resulted in GLP-1 secretion, and these bacterial acyl dopamines were more potent than OEA, free fatty acids, and dopamine (Supplemental Figure 8). Acyl dopamines did not induce GLP-1 secretion in the non-differentiated (non-EEC-containing) intestinal organoids (Figure 4F).

### *Blautia wexlerae* and acyl amines induced GLP-1 secretion in mice and improved glycemic and appetite control

We next sought to determine if *B. wexlerae* and its acyl amine metabolites may influence metabolic outcomes by EEC activation in mice. As *B. wexlerae* could not successfully mono-colonize germ-free mice, per absent bacteria on gram stain of stool and no recovery of colonies from stool cultures, we co-colonized germ-free mice with *Bacteroides thetaiotamicron* (*B. theta*) and *B. wexlerae* together, using the *B. theta* partner to condition the gut environment for *B. wexlerae*. This 2-plex consortium (Figure 5A) enabled successful engraftment of germ-free mice with *B. wexlerae,* as evidenced by recovery of live *B. theta* and *B. wexlerae* from stool samples. Stool metabolomics revealed a distinct metabolomic profile in mice colonized with *B. theta + B. wexlerae* compared to germ-free mice or mice colonized with *B. theta* alone (Figure 5B, Supplemental Table 3). Acyl amines were among the top 20 differentially abundant stool metabolites in *B. theta + B. wexlerae* colonized mice (Supplemental Figure 9, Supplemental Table 4). Specifically, acyl ethanolamines were significantly more abundant in the *B. theta + B. wexlerae* co-colonized mice (Figure 5C), and acyl dopamine levels trended higher in both colonization groups, though this was not statistically significant (Supplemental Figure 10).

**Figure 5.**
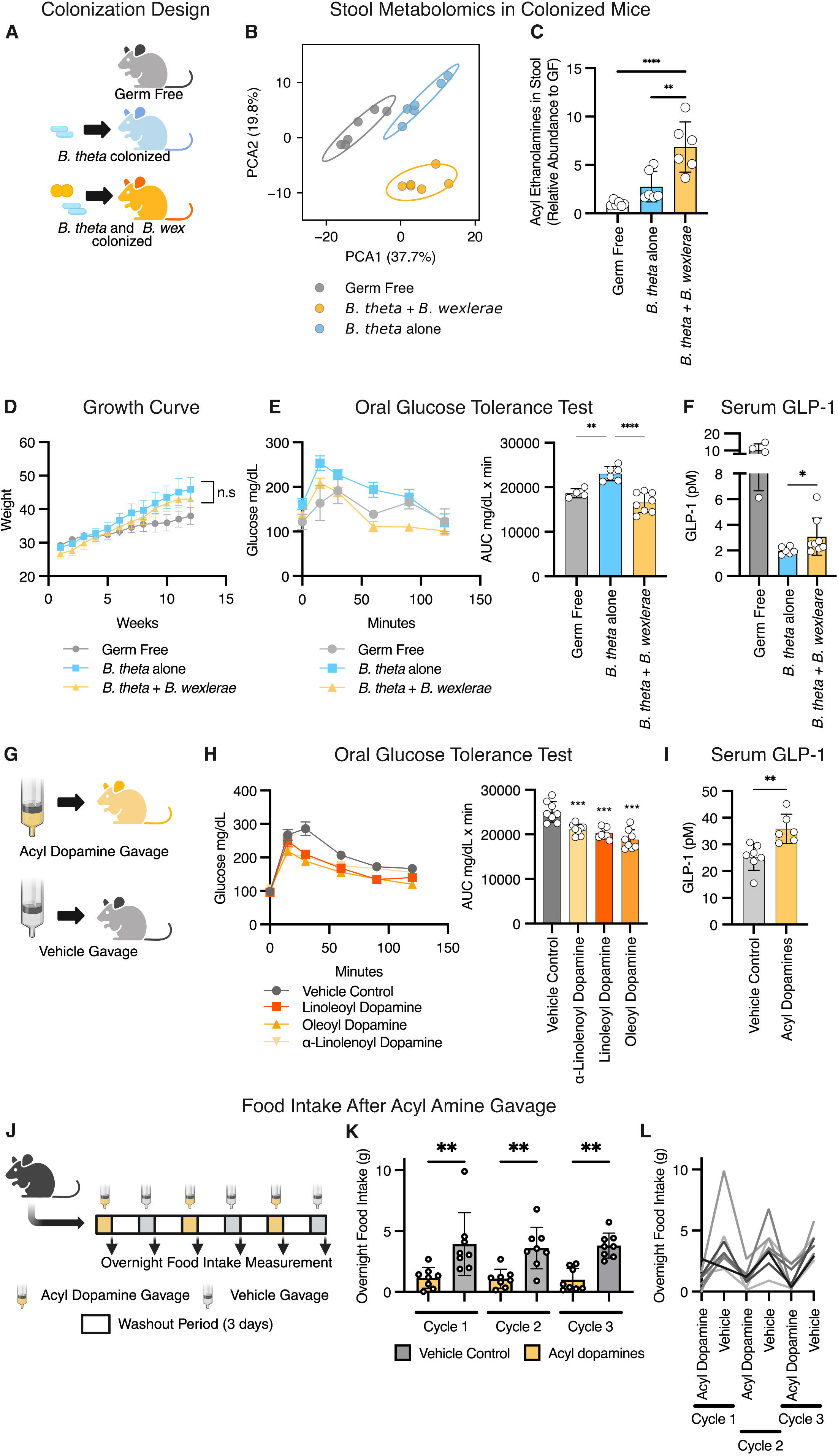
Blautia wexlerae and acyl amines induce GLP-1 secretion in mice and improve glycemic control. **A.** Schematic of the three groups of colonized mice: germ-free mice, germ-free mice mono-colonized with *Bacteroides thetaiotamicron*, and germ-free mice colonized with both *B. theta* and *Blautia wexlerae*. **B.** PCA plot from stool LC-MS metabolomic analysis, showing distinct metabolomes between the three groups. **C.** Relative abundance (reference group: germ-free) of acyl ethanolamines, quantified from the stool by LC-MS. Mice colonized with both *B. theta* and *Blautia wexlerae* had the highest abundance of stool acyl ethanolamines. **D.** Growth curves in all three groups of male mice placed on a high-fat (45% fat) diet. Germ-free mice are resistant to diet-induced obesity, and there was no difference in weight gain between the *B. theta* alone and *B. theta + B. wexlerae* colonized mice. **E.** Oral glucose tolerance test (2 mg/kg) in mice 30 minutes with area under the curve analysis. *B. theta + B. wexlerae* colonized mice exhibited improved glycemic control. **F.** GLP-1 secretion in plasma sampled 20 minutes after glucose gavage (2 g/kg). *B. theta + B. wexlerae* colonized mice had increased GLP-1 secretion compared to *B. theta* alone. **G.** Schematic of the acyl amine gavage experiments. **H.** Oral glucose tolerance test (2 mg/kg) in mice 30 minutes after treatment via gavage with *Blautia* acyl dopamines (0.1 g/kg). Area under the curve analysis shown on the right. All three acyl dopamines improved glycemic control. **I.** GLP-1 secretion in plasma sampled 20 minutes after gavage with a combination of three acyl dopamines (0.1 g/kg total) followed immediately by a glucose gavage (2 g/kg). **J.** Schematic of food intake experiment. **K.** Mean and distribution of overnight food intake after gavage with a combination of three acyl dopamines (0.1 g/kg total) vs. vehicle. **L.** Individually traced overnight food intake, across three cycles of acyl dopamine or vehicle gavage; each line represents a single mouse. (*, *p* < 0.05; **, *p* < 0.01; ***, *p* < 0.001; ****, *p* < 0.0001).

We next put germ-free mice colonized with *B. theta* alone or mice with *B. theta + B. wexlerae* together on a high-fat diet (45% fat) rich in MUFAs and PUFAs. As expected, germ-free mice showed resistance to diet-induced obesity (Figure 5D). In male mice, we found no difference in weight gain between the *B. theta* alone and *B. theta + B. wexlerae* colonized groups (Figure 5D). However, the male mice colonized with *B. theta* + *B. wexlerae* exhibited a significant improvement in glycemic control compared to mice colonized with *B. theta* alone (Figure 5E). Male mice colonized with *B. theta + B. wexlerae* also showed a stronger GLP-1 response, implicating an effect of *B. wexlerae* on EECs and GLP-1 secretion (Figure 5F and Supplemental Figure 9). As expected, the obesity-resistant GF mice used as controls had higher GLP-1 secretion overall (Figure 5F). After glucose gavage, male mice colonized with *B. theta + B. wexlerae* also exhibited higher circulating levels of another EEC secreted gut hormone, glucose-dependent insulinotropic peptide (GIP), compared to mice colonized with *B. theta* alone (Supplemental Figure 10). In contrast, and consistent with sexually dimorphic metabolic and enteroendocrine outcomes^33^, female mice showed a relative resistance to diet-induced obesity and no detectable differences in weight or metabolic outcomes regardless of gut colonization with *B. theta* alone or *B. theta + B. wexlerae* together (Supplemental Figure 10).

Given the strong induction of GLP-1 secretion in culture models and EEC activation in male mice, we hypothesized that the three most potent *Blautia*-produced acyl dopamines might improve glycemic control when fed to mice in vivo. To test this hypothesis, we administered oleoyl-, linoleoyl-, and α-linolenoyl-dopamine (0.1 g/kg by gastric gavage) and performed an oral glucose tolerance test (Figure 5G). We found that all three compounds significantly improved glycemic control following a single dose (Figure 5H). We then tested a mixture of all three acyl dopamines (total dose of 0.1 g/kg) and measured GLP-1 secretion directly. Acyl dopamine-treated mice exhibited significantly higher GLP-1 secretion 20 minutes after compound and glucose gavage (Figure 5I). We then treated mice with a combination of the three acyl dopamines (oleoyl-, linoleoyl-, and α-linolenoyl-dopamine) and measured food intake (Figure 5J).

Consistent with an agonist effect on gut EECs, we found that mice gavaged with the acyl dopamines showed a nearly 3-fold reduction in food intake (Figure 5K and 5L).

These studies demonstrate that *B. wexlerae* colonization or acyl dopamine treatment improve post-prandial glycemic control and gut peptide hormone responses in vivo, thus defining a mechanism of action for how *B. wexlerae* can affect metabolic health^34,35^.

### Acyl amine synthesis correlates with leanness and decreased fatty intake in humans

The association between *B. wexlerae* colonization and leanness in humans has been inconsistently observed, even in large multicenter studies^34–39^. Given our results, we hypothesized that measurement of *Blautia*-derived acyl amine production, rather than *Blautia* stool abundance, may link more strongly to lower body weight in humans. To explore this, we aligned human shotgun metagenomic sequences from the Global Microbiome Conservancy project with the *B. wexlerae* CTA operon (Figure 6A). Using an alignment cutoff of 90%, we calculated the number of aligned reads per million total reads for each metagenome. We found that individuals with normal weights (BMI 18.5 to 25 kg/m²) had higher amounts of *B. wexlerae* specific CTA-mapped reads compared to individuals with obesity (BMI ≥ 30 kg/m²) (Figure 6B). Notably, stool content of *B. wexlerae* itself did not directly correlate with leanness in this cohort^40^. When we expanded our analysis to include homologs from other *Lachnospiraceae* species, the association with leanness also did not hold (Supplemental Figure 11). These results suggest that only the acyl amine synthesis genes specific to *Blautia* link to a healthy weight in humans (Supplemental Figure 11). A similar trend was observed in healthy children from the NIH cohort, who had higher relative *Blautia*-specific CTA-aligning reads compared to adolescents from the same study with severe obesity (Figure 6C).

**Figure 6.**
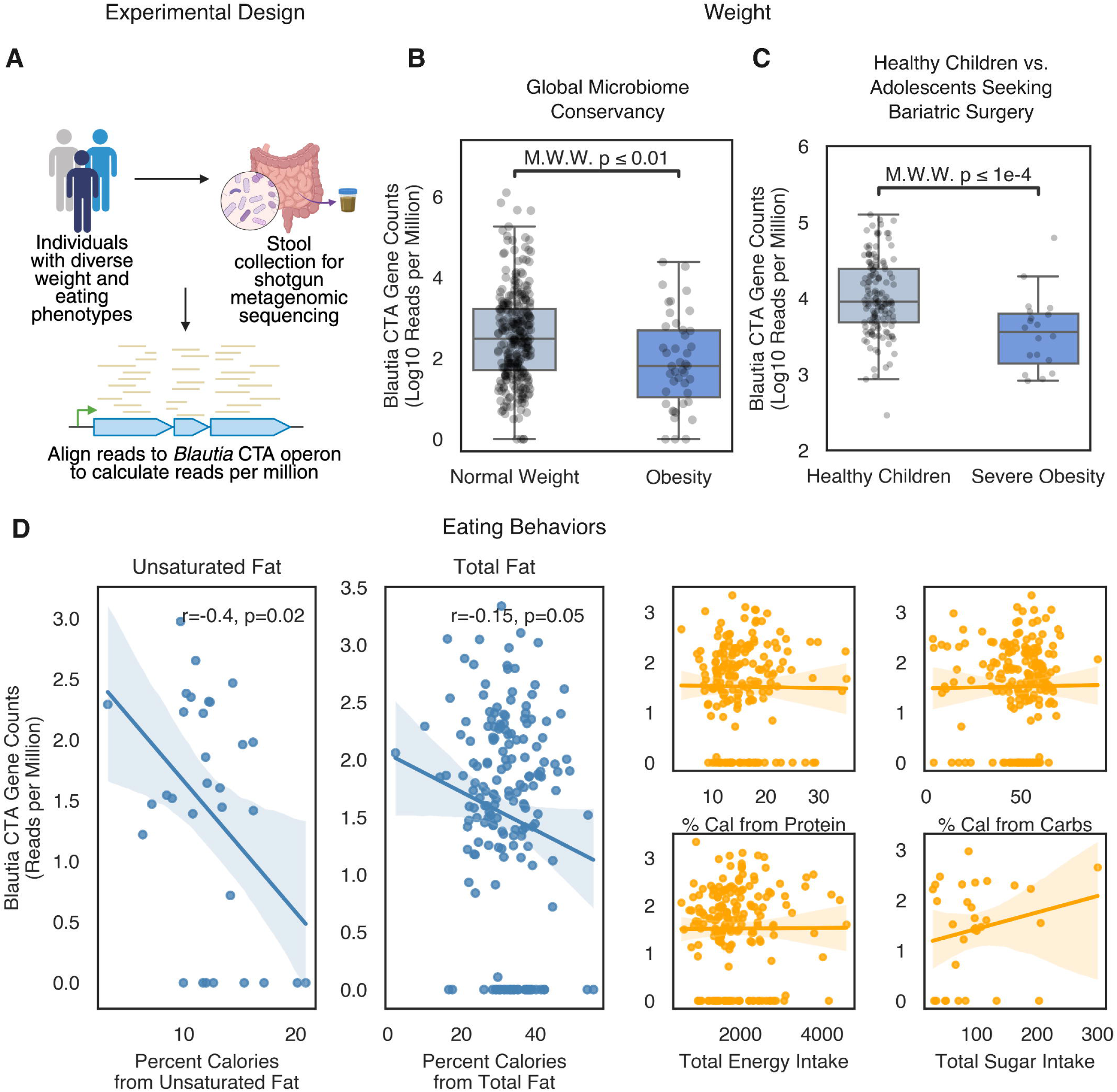
Blautia acyl amine genes are associated with leanness and decreased fatty acid intake in humans. **A.** Schematic of metagenomic sequencing analysis with direct mapping of reads to *Blautia* or *Lachnospiraceae* CTA cassette sequences. **B.** Comparison between individuals with normal weight vs. obesity in the Global Microbiome Conservancy. *Blautia* acyl amine gene abundance was higher in normal weighted individuals. **C.** Comparison between healthy children and adolescents seeking bariatric surgery. *Blautia* acyl amine gene abundance was higher in healthy children. **D.** Correlations of *Blautia* acyl amine gene abundance with fat, unsaturated fat, protein, carbohydrate and total caloric intake in children. *Blautia* acyl amine gene abundance was negatively correlated with both fat and unsaturated fat intake (r and p-values calculated using Spearman’s correlation).

These results predict that acyl amine synthesis by *Blautia* could be associated with healthier eating habits in humans. In accord with this prediction, we found a greater abundance of microbial DNA encoding *Blautia-specific* acyl amine synthesis genes in the stools of individuals who did not report LOC eating (Supplemental Figure 12). Mechanistically, we reasoned that such bacterially produced acyl amines, which require dietary fatty acids as substrates, might specifically reduce the host’s preference for dietary fats by enhancing fatty acid-driven satiety-producing gut hormone release, a response already known to influence eating habits and food preferences through reward circuitry^41^. To test this, we conducted dietitian-administered dietary recalls in healthy children and measured the relative intake of fat, protein, and carbohydrates. We found that children with higher stool levels of *Blautia* acyl amine synthesis genes consumed less total fat and unsaturated fat compared to children with lower levels of acyl amine synthesis genes in their stool (Figure 6D). Relative amounts of protein and carbohydrates, as well as total ingested calories, were similar across both cohorts.

## DISCUSSION

*Blautia wexlerae,* a common gut commensal, encodes a gene operon that can synthesize acyl amines from dietary fats. These bacterial acyl amines act as hormone secretagogues, inducing GLP-1 secretion with greater potency than endogenously produced acyl amines in vitro. In vivo, the bacterial acyl amines activate gut enteroendocrine cells to secrete GLP-1, improve glucose metabolism, and decrease food intake, thus elucidating a microbial mechanism for modulating host endocrine signaling.

Our findings align with growing, albeit mixed, evidence that *Blautia* species support metabolic health in humans^42–45^. We followed three independent human cohorts, all of which demonstrated a positive association between *Blautia* colonization and low weight and/or healthier eating behaviors. Mexican children with obesity, especially those with insulin resistance, have lower levels of *B. wexlerae* in their stool samples compared to healthy controls^37^. Similarly, Japanese adults with obesity exhibit relative depletion of *B. wexlerae*^36^. The association between diminished *B. wexlerae* colonization and LOC eating in our study also resonates with Samulenaite and colleagues’ recent report of lower *Blautia* stool levels in humans with food addiction^34^. The mechanism of action defined by the studies reported here helps to explain these emerging associations between *Blautia* abundance and metabolic health.

We also found a fitness advantage that *Blautia* gains from acyl amine production and resistance to free fatty acids, thus providing a co-evolutionary hypothesis for why *Blautia’s* acquisition of the CTA operon. High concentrations of free fatty acids, comparable to those seen in the gut lumen in vivo, were found toxic to *Blautia* and related *Lachnospiraceae*. This discovery aligned with a recent report that free fatty acids were toxic to *Blautia producta* and *Blautia luti*^46^. Remarkably, the fatty acid toxicity on *Blautia* growth in vitro was mitigated by the addition of free amines. Thus, acyl amine synthesis likely evolved to improve microbial fitness within the gut environment and has been secondarily co-opted by the host for its beneficial metabolic effects.

From the host perspective, microbial acyl amines appear to act as potent EEC peptide hormone secretagogues. Other compounds with similar EEC-agonist activity have had mixed effects when applied in human trials^47^. But the newer generations of such agonists, including recent GPR40 and GPR119 agonists, have shown improved glycemic control and weight loss in clinical trials^48,49^. Because microbes such as *Blautia* colonizing the gut have direct access to the enteroendocrine system^11,12^, and secrete potent acyl amine GPR agonists, this pathway of host-microbial cross-talk may show similar efficacy in affecting EEC cells and host metabolism. Though selectively inhibiting GLP-1-producing EECs in the mouse gut has not always resulted in hyperphagia or weight gain (suggesting a compensatory mechanism in these models^14,15^), recent studies using intersectional genetics have demonstrated that stimulating various EEC subtypes, such as GLP-1-producing ileal and colonic L-cells, GIP-producing K-cells, and enterochromaffin cells, can effectively induce satiety^16–19^. Notably, *Blautia* species are enriched in these EEC-rich regions of the distal GI tract, perhaps contributing decisively to the actions of *Blautia*-derived metabolites^11,50,51^.

While other microbial metabolites can induce GLP-1 secretion, several factors point to *Blautia* acyl amines as important drivers in this pathway. Unlike other microbes implicated in GLP-1 induction, such as *Gemella*, *Blautia* are commonly found and abundant in the human distal GI tract^52^. Furthermore, the acyl dopamines produced are more potent EEC activators than other proposed microbial ligands^52–56^. Regarding other possible active *B. wexlerae* metabolites, previous studies in mice implicated the production of short-chain fatty acid (SCFA) by *B. wexlerae* as a mechanism of action^35^. However, the results of those studies were confounded by the lack of demonstrable *B. wexlerae* colonization^34,35^, and the increase in stool SCFAs induced by *Blautia* treatment could have originated from other gut commensals^35^. Lastly, acyl amines have also been implicated in other pathways affecting the host. Dohnalova et al., for example, recently demonstrated in mice that microbiome-driven increases in stool acyl amines activated TRPV1 receptors on gut sensory neurons and promoted dopamine signaling during exercise, thus leading to increased physical activity^57^.

In summary, we identify *Blautia wexlerae* as a key microbe linked to LOC eating in children and define a mechanism through which *B. wexlerae* contributes to satiety signaling in mice and likely humans. GLP-1 receptor agonists have revolutionized obesity therapy, but they remain limited by cost, tolerance, and durability of effect. In the real world, outside of clinical trials, upwards of 70% of patients stop using these drugs at two years^58–60^. Thus, there remains strong motivation for more natural microbiome-based approaches to treat obesity and metabolic disorders, like those suggested by *Akkermansia muciniphila*^61^. Such an approach to harness the gut microbiome as treatment for obesity may prove to have longer-lasting effects and to be more appropriate for obesity prevention. Our findings provide rationale for engineering the human microbiome to boost *B. wexlerae* abundance and its production of acyl amines as treatment for disordered eating behaviors and obesity.

## METHODS

### Participant Recruitment

NIH. Starting in 2015, generally healthy participants aged 8-27 years from Maryland, Virginia, and the District of Columbia were recruited for the Children’s Growth and Behavior study (NCT02390765), an ongoing, prospective longitudinal study of eating behavior, by mailed advertisements and posted flyers in neighborhoods within 50 miles of the NIH Clinical Center (Bethesda, MD). Youth were excluded from the parent study if they were underweight (BMI < 5th percentile), had experienced recent weight loss (> 5% of body weight), were regularly using medications or substances known to impact weight and/or eating behaviors, had a history of significant or recent brain injury, had a Full Scale Intelligent Quotient ≤ 70^62^, or a diagnosis of a serious medical condition or full threshold Diagnostic and Statistical Manual 5th Edition^63^ disorder, other than binge-eating disorder. Exclusionary psychiatric conditions were assessed by the Schedule for Affective Disorders and Schizophrenia for School- Age Children^64^ and the Eating Disorder Examination (child or adult version)^65,66^, respectively. All eligible participants were offered the opportunity to supply stool samples as an option for which they were given additional compensation, but stool collections were not required for participation. Before any study procedures were administered, informed consent from parents and/or guardians and assent from the children were obtained. Study procedures were approved by the National Institutes of Health institutional review board.

From 2016 to 2021, participants aged 14 to 26 were recruited from the Adolescent Bariatric Surgery Program at Boston Children’s Hospital (BCH). This study was approved by the Institutional Review Board at Boston Children’s Hospital. All participants were eligible for bariatric surgery using criteria set by the American Board of Metabolic and Bariatric Surgery. However, sampling was significantly interrupted during the COVID-19 pandemic, so only 6 participants underwent surgery during the study. These participants also submitted stool samples and eating behavior assessments at 3 and 12 months postoperatively.

### Eating Behavior Assessments

LOC eating. Participants underwent semi-structured interviews, either The Eating Disorder Examination adult^66^ or child^65^ version, to assess LOC eating within the past one month. The Eating Disorder Examination contains 21 items that assess disordered attitudes and behaviors related to eating, body-shape and weight, and 13 items^66^ designed and adapted to diagnose specific DSM-5 eating disorders^63^. The Eating Disorder Examination has demonstrated sound psychometric properties, including good to excellent test-retest and interrater reliability in youth across the weight spectrum, as well as individuals from diverse racial/ethnic identity backgrounds^67–70^. The child version of Eating Disorder Examination has demonstrated good to excellent interrater reliability and good internal consistency and discriminant validity among youth^65,71,72^. Training and administration for the Eating Disorder Examination was performed as previously described^71^. Participants in the BCH Bariatric Surgery clinic were administered the Yale Food Addiction Scale, which includes a loss of control subscale^22,73^.

Dietary recall. Participants, with the assistance of their parents/guardians, were interviewed by research dietitians, using a multi-pass approach to limit the extent of under-reporting, to obtain information on intake during the 24 hours before their visit. This method has been validated against the doubly labeled water method in children^74^.

### Stool samples and sequencing analysis

Stool samples were collected at home by participants at the National Institutes of Health (NIH), transported on ice, and stored at -80°C. For participants at Boston Children’s Hospital (BCH), stool samples were collected using a wipe-based protocol, as previously described, and stored at -80°C. DNA was extracted from all samples (NIH and BCH) in a single batch using the PowerSoil DNA Isolation Kit (Qiagen). Shotgun metagenomic sequencing was performed by the Microbial ‘Omics Core at the Broad Institute (Cambridge, Massachusetts). Sequence quality was assessed using FastQC, and taxonomic classification was conducted using Kraken2 and Bracken, which mapped the data to a custom Kraken database containing newly sequenced genomes from the Global Microbiome Conservancy^75,76^.

### Machine Learning

To distinguish the microbiomes of participants with and without Loss of Control (LOC) eating, we employed strategies to address the imbalance in the minority LOC eating class. Initially, the dataset was split into training (75%) and test (25%) sets. To mitigate class imbalance, the Synthetic Minority Oversampling Technique (SMOTE) from the imbalanced-learn package (imblearn) was applied to increase the LOC eating class, ensuring that it comprised 25% of the total training set^77^. A random forest classifier (scikit-learn version 1.6.1) was then trained to differentiate between the two groups.

Additionally, a second random forest classifier was trained on a modified training set, where the LOC class was oversampled to 25% of the total training set, to further investigate model performance under different class distributions.

### Differential Abundance Testing

Relative abundance at the species level was used to identify taxonomic differences between participants with and without Loss of Control (LOC) eating. Species relative abundances were calculated using the Kraken2/Bracken pipeline, as described above. Species identified in fewer than 20% of stool samples were excluded from the analysis. Differential relative abundance between the two groups for each species was assessed using the non-parametric Mann-Whitney U test, with multiple hypothesis testing corrections applied using the Benjamini-Hochberg method^78^. Species were considered differentially abundant if the false discovery rate (FDR) was less than 0.05. In the bariatric surgery cohort, the differential relative abundance of *Blautia wexlerae* at baseline was specifically tested using the Mann-Whitney U test to assess statistical significance.

### Homology Search in the Global Microbiome Conservancy

A protein-sequence BLAST database was constructed using genomes from the Global Microbiome Conservancy (GMBC) and the Broad Institute-OpenBiome Microbiome Library (BIO-ML) to identify homologs of the CTA cassette^28,29^. The database included a total of 7,781 genomes. The search queries targeted all acyl transferase proteins (C proteins) and fatty acyl-CoA ligase proteins (A proteins) from Clostridia, as characterized and cataloged in the Chang et al. manuscript^30^. The acyl carrier protein (T protein), being much smaller, was excluded from the homology search. An isolate was considered to have a CTA cassette homolog if it encoded both C and A gene homologs (using cutoffs of 70% amino acid identity across 80% of the protein sequence) and if these genes were located within a putative operon structure. Notably, all isolates with identified homologs exhibited the C and A genes in an operonic structure with an acyl carrier (T) protein situated between them. Species identification and phylogenetic reconstructions were conducted as described previously^28,29^. For each species cluster, the presence or absence of CTA cassette homologs was recorded by determining the number of isolates with and without homologs.

### Metagenomic alignments using Diamond

To evaluate the relative abundance of acyl amine synthesis gene clusters in human metagenomic samples, we employed the protein sequence aligner Diamond^79^. We constructed two separate databases: one consisting of *Blautia*-specific CTA cassette homologs identified in the GMbC and BIO-ML isolates, and another containing CTA cassette homologs from all Clostridia within the same isolates. Notably, no homologs were found outside of the Clostridia class. Each shotgun metagenomic read from participant stool samples was aligned using Diamond. A read was considered to align with a *Blautia* or Clostridia CTA homolog if it exhibited at least 90% homology over a 30-amino acid region to one of the CTA genes in the databases. For each sample, we calculated the counts of aligned reads per million total reads, using this metric as a measure of the relative abundance of *Blautia*-specific or Clostridia acyl amine synthesis genes. In the GMbC cohort, we compared the acyl amine reads per million between participants with normal weight (BMI 18.5–25 kg/m²) and those with obesity (BMI ≥25 kg/m²), testing for statistical significance using the Mann-Whitney U test. Additionally, we examined the association between acyl amine reads per million and food intake metrics, employing Spearman’s rank correlation test for statistical analysis.

### Pathway enrichment in Blautia using a TwinsUK dataset

Microbe-metabolome associations and represented pathways were downloaded from supplemental tables from Zierer et al. report from the TwinsUK study^24^ . For pathways enrichment with *Blautia*, we evaluated over-representation of the predefined metabolic pathways using the hypergeometric distribution. N was defined as the total number of metabolites in the annotated dataset, K the number of metabolites significantly associated with Blautia, n the number of metabolites belonging to the given sub-pathway in the full dataset, and k the number of Blautia-associated metabolites within that sub-pathway. P-values were calculated in Python (SciPy) using the survival function of the hypergeometric distribution, with false-discovery-rate (FDR) calculated post hoc using the Benjamini-Hochberg method.

### Culture conditions and fatty acid supplementation

Individual isolates from the GMbC and BIO-ML collections were revived from frozen glycerol stocks by plating onto Brucella Blood Agar plates (Thermo Fisher; Waltham, MA) in an anaerobic chamber (Coy; Grass Lake, MI), and their taxonomy was confirmed through 16S sequencing. Colonies were then inoculated into either Reinforced Clostridial Media (RCM, Sigma-Aldrich; St. Louis, MO) or de Man, Rogosa, and Sharpe medium (MRS, Sigma-Aldrich; St. Louis, MO) under anaerobic conditions. To assess the effect of free fatty acids on bacterial growth, liquid media was supplemented with 100 μM of oleic acid, linoleic acid, or α-linolenic acid (Sigma-Aldrich; St. Louis, MO), vortexed for 5 minutes to ensure thorough mixing, and incubated in the anaerobic chamber for at least 24 hours before use. Cultures were grown in 1 mL volumes in 96-well deep-well plates (Thermo Fisher; Waltham, MA) at 37°C, with each experimental condition tested in triplicate. Optical density (OD600) was measured after 24 to 36 hours of incubation. The relative growth in high-fat conditions was calculated by dividing the OD600 of each triplicate by the average OD600 of cultures grown in non-supplemented media.

### LC-MS for untargeted metabolomics and acyl amine detection

Metabolites were profiled using an LC–MS system consisting of a Nexera X2 U-HPLC (Shimadzu Scientific Instruments; Marlborough, MA) coupled to an IDX orbitrap mass spectrometer (Thermo Fisher Scientific; Waltham, MA). Cell pellets were resuspended in 5 volumes of 90% methanol and media (30 ul) extracted using 90 μL methanol. For stool metabolite analysis from both human and mouse, frozen stool samples were homogenized in 10 μl of water per mg of stool weight using a bead mill (TissueLyser II; Qiagen) for 4 minutes. The aqueous homogenates were aliquoted for metabolite profiling analyses. For bacterial cell extracts, samples were vortexed for 1 minute at room temperature and centrifuged (10 min, 15,000g, 4°C). The supernatants (10 μL) were injected directly onto a Acquity UPLC HSS T3 Column; 1.8µm, 2.1X150mm (P/N 186003540 Waters; Milford, MA). The column was eluted at a flow rate of 400 µl/min with initial conditions of 5% mobile phase B (0.1% Formic Acid in Acetonitrile) and 95% mobile phase A (0.1% Formic Acid in Water) for 3 minutes, followed by a 17 min linear gradient to 100% mobile phase B. Mass spectrometry analyses were carried out using electrospray ionization in the positive ion mode using full scan analysis over *m*/*z* 80–1000 at 120,000 resolution. Additional mass spectrometry settings were: ion spray voltage, 3.5 kV; ion transfer temperature, 320 °C; vaporizer temperature, 300 °C; sheath gas, 40; auxiliary gas, 8; and S-lens RF level % 70.

Raw data were processed using Progenesis QI software version 2.0 (NonLinear Dynamics) for feature alignment, nontargeted signal detection, and signal integration. Targeted processing of oleoyl dopamine was conducted using TraceFinder software version 4.1 (Thermo Fisher Scientific; Waltham, MA). The compound identities were confirmed using reference standards and to elucidate potential acyl amine matches in the data we generated an *in silico* combinatorial database of 460 amino-containing compounds and 53 fatty acids for a total of 24,380 acyl-amines and 7,578 unique formulas. Using the predicted *m/z* for these compounds we searched for matches in the non-targeted data at 5 ppm mass accuracy. Unknown features with potential acyl-amine matches were targeted for tandem MS (MS/MS) using higher-energy collisional dissociation (HCD) at 20,30,40, and 50 eV HCD. The resulting fragmentation patterns for each unknown were used to screen for diagnostic fragments for the in silico matched acyl amine at a 10 ppm mass accuracy window. For instance, for Palmitoyl Arginine (C22H44N4O3), we monitored the diagnostic fragment for the amine-containing moiety (Arginine [C6H14N4O2+H]^+^ m/z 175.1189) and the amine-containing fatty acid fragment (palmitamide [C16H33NO+H]^+^ m/z 283.2631).

### Screening for amines that rescue growth in Blautia

*Blautia wexlerae* (strain 59E11 from the GMbC) was cultivated in Reinforced Clostridial Media (RCM) alone or in RCM supplemented with 100 μM free fatty acids, as described above. Individual colonies were initially inoculated in RCM, then back-diluted at a 1:100 ratio twice prior to initiating the screen. To identify amines that could rescue the growth of *Blautia wexlerae* under high-fat conditions, a series of amines (see the Results section for a list of amines tested) were spiked into the media at a concentration of 100 μM. Each condition was tested in triplicate. All cultures were grown in 1 mL volumes in 96-well deep-well plates at 37°C in an anaerobic chamber. At 16, 20, and 24 hours post-inoculation, a small aliquot was removed from each culture to measure optical density (OD). At the time points when growth in RCM alone reached its maximum (OD600 of 0.6–0.8), and when growth in RCM supplemented with free fatty acids was suppressed (OD600 of 0.1–0.2), the relative rescue effect of the amines was calculated by determining the OD600 ratio of amine-supplemented vs. non-amine-supplemented high-fat media. To validate these findings, we tested an extended dilution series (1, 10, and 100 μM) of both fatty acids and amines in triplicate by generating growth curves in a 96-well plate format (200 μL per well) and measuring OD using the Stratus kinetic and endpoint microplate reader (Cerillo; Charlottesville, VA). Additionally, the growth of *Blautia* with 100 μM acyl amine supplementation was assessed using the microplate reader and analyzed on GraphPad Prism 10.

### GPR119 assay

We generated a line of GPR119 bioreporter cells by transfecting HEK293T cells, which express a CRE-luciferase reporter construct (BPS Bioscience; San Diego, CA), with a plasmid overexpressing the human GPR119 open reading frame (OriGene; Rockville, MD). The system was validated using the small molecule GPR119 agonist AR231453^80^ and the endogenous ligand oleoyl ethanolamine (OEA)^31^ . The EC50 values for both compounds were consistent with those reported for other luciferase or cyclic AMP-based reporter systems. Acyl amines, identified through human metabolomics and an *ex vivo* screen using Clostridial acyl transferases, were tested on this bioreporter cell line. Except for Oleoyl Dopamine and OEA (Sigma-Aldrich; St. Louis, MO), the acyl amines were synthesized as below and confirmed by GC-MS. Acyl amines were dissolved in Krebs buffer with 1% DMSO and tested at a top concentration of 100 μM, with an 8- to 12-point serial dilution series used to determine the EC50 for each compound (GraphPad Prism 10). Bioreporter cells were incubated with the acyl amines for 2–3 hours, and GPR119 activity was measured by assessing luciferase signal using the Bright-Glo Luciferase Reagent (Promega; Madison, WI) using standard kit conditions and protocols. We also tested each compound on mock-transfected CRE-luciferase cells to demonstrate that there was no non-specific activation of the CRE-luciferase system by any of the compounds.

### Acyl dopamine synthesis

All solvents and reagents were used directly from commercial suppliers. Analytical TLC was performed on silica gel 60/F254 pre-coated aluminum sheets (0.25 mm, Merck) and visualized using KMnO_4_ TLC stain. Flash column chromatography purification was performed using silica gel 60 (0.04–0.063 mm, 230–400 mesh) and HPLC grade solvents. Low-resolution mass spectrometry (LRMS) analysis: Agilent 1200 system equipped with G1379B degasser, G1312A binary pump, G1329A autosampler, G1316A thermostatted column compartment (TCC), G1315D DAD, and 6130 quadrupole LC/MS. ^1^H Nuclear Magnetic Resonance (NMR) spectra were conducted on Varian 400 MHz and/or Bruker Avance 400 MHz spectrometer, and obtained at a frequency indicated in detailed experimental below. The chemical shifts of ^1^H spectra are reported in parts per million (ppm). For all ^1^H NMRs, the chemical shifts of all NMR were measured relative to the expected solvent peaks of the respective NMR solvents: CDCl_3_ (7.26 ppm). The data for all spectra is reported in the following format: chemical shift (multiplicity, coupling constant, integration). Multiplicity is defined by examples: s = singlet, d = singlet, d = doublet, t = triplet, q = quartet, sd = singlet of doublets, dd = doublet of doublets, dt = doublet of triplets, tt = triplet of triplets, ddd = doublet of doublet of doublets, and m = multiplet. A broad resonance is denoted by the abbreviation br and the apparent splitting is denoted as the abbreviation app. Coupling constants are applied as J in Hertz (Hz).

Oleoyl Dopamine Synthesis. Step 1: To a solution of oleic acid (3.0 g, 10.62 mmol) in dry DCM (10 mL) under argon was slowly added oxalyl chloride (4.05 g/ 2.73 mL, 31.91 mmol) over 15-20 min. The resulting mixture was stirred at room temperature overnight then concentrated *in vacuo* to give the crude oleoyl chloride which was used directly in the next step without further purification. Step 2: A solution of dopamine HCl salt (2.42 g, 12.76 mmol) in dry DMF (45 mL) was cooled to 0 °C under argon. Dry triethylamine (4.3 g/ 5.93 mL, 42.49 mmol) was added and the mixture was stirred at this temperature for 5 min. To this mixture was then added dropwise a solution of the crude oleoyl chloride from the last step in dry DMF (15 mL). The resulting mixture was protected from light, stirred at 0 °C for 1.5 hr then room temperature overnight. The reaction mixture was then diluted with diethyl ether, washed with 1N HCl x1, 0.5N HCl x2, dried over Na_2_SO_4_ and concentrated *in vacuo*. The resulting crude residue was purified using flash column chromatography (0 – 50% ethyl acetate in hexanes + 1% acetic acid). Fractions containing the desired product were combined and concentrated. The residue was re-dissolved in diethyl ether (50 mL), washed with 2% K_2_CO_3_ solution x1 (20 mL), 1N HCl x1 (20 mL), dried over Na_2_SO_4_ and concentrated *in vacuo* (water bath temp ∼10 °C). The resulting residue was then triturated with cold hexanes and filtered to give the title compound as a white solid (3.42g, 77%) which was further dried under high vac at 4 °C. Estimated melting point: > 25 °C. ^1^H NMR (400 MHz, CDCl_3_) δ 7.41 (s, 1H), 6.81 (d, *J* = 8.0 Hz, 1H), 6.75 (sd, 1H), 6.57 (dd, *J* = 8.1, 2.1 Hz, 1H), 5.81 (s, 1H), 5.57 (br t, *J* = 5.9 Hz, 1H), 5.40 – 5.27 (m, 2H), 3.49 (td, *J* = 7.1, 5.8 Hz, 2H), 2.70 (t, *J* = 7.1 Hz, 2H), 2.15 (app. t, 2H), 2.05 – 1.96 (m, 4H), 1.63 – 1.58 (m, 2H, partially overlapped with water peak), 1.35 – 1.25 (m, 20H), 0.88 (app. t, 3H). MS (ESI^+^): 418.3.

*N*-Linoleoyl dopamine was synthesized in analogy to that of *N*-oleoyl dopamine. Starting with 2.5 g of linoleic acid, *N*-linoleoyl dopamine was obtained as a white solid (2.78 g, 75%). Estimated melting point: 15 – 20 °C. ^1^H NMR (400 MHz, CDCl_3_) δ 7.44 (br s, 1H), 6.81 (d, *J* = 8.0 Hz, 1H), 6.75 (sd, *J* = 2.1 Hz, 1H), 6.57 (dd, *J* = 8.0, 2.1 Hz, 1H), 5.83 (br s, 1H), 5.58 (br t, *J* = 5.9 Hz, 1H), 5.43 – 5.27 (m, 4H), 3.49 (td, *J* = 7.1, 5.8 Hz, 2H), 2.77 (app. t, 2H), 2.70 (t, *J* = 7.1 Hz, 2H), 2.15 (app. t, 2H), 2.09 – 1.99 (m, 4H), 1.61 – 1.56 (m, 2H, partially overlapped with water peak), 1.39 – 1.25 (m, 14H), 0.89 (app. t, 3H). MS (ESI^+^): 416.3.

*N*-α-Linolenoyl dopamine was synthesized in analogy to that of *N*-oleoyl dopamine. Starting with 3.0 g of α-linolenic acid, *N*-α-linolenoyl dopamine was obtained as a white solid (3.75 g, 84%). Estimated melting point: 10 – 15 °C. ^1^H NMR (400 MHz, CDCl_3_) δ 7.49 (s, 1H), 6.80 (d, *J* = 8.0 Hz, 1H), 6.75 (sd, *J* = 2.1 Hz, 1H), 6.57 (dd, *J* = 8.1, 2.1 Hz, 1H), 5.87 (s, 1H), 5.58 (br t, *J* = 5.7 Hz, 1H), 5.45 – 5.26 (m, 6H), 3.49 (td, *J* = 7.1, 5.9 Hz, 2H), 2.85 – 2.76 (m, 4H), 2.70 (t, *J* = 7.1 Hz, 2H), 2.15 (app. t, 2H), 2.12 – 2.00 (m, 4H), 1.59 – 1.53 (m, 2H, partially overlapped with water peak), 1.36 – 1.25 (m, 8H), 0.97 (t, *J* = 7.5 Hz, 3H). MS (ESI^+^): 414.3.

### NCI cell assay and GLP-1 measurement

To assess the ability of acyl amines to induce GLP-1 secretion, we utilized the enteroendocrine cell line NCI-h716. NCI-h716 cells were plated in 96-well plates on Matrigel (Corning; Corning, NY) and incubated for two days. Following this, cells were washed with PBS, starved in PBS for 30 minutes, and treated with serial concentrations (100, 50, 25, and 12.5 μM) of acyl amines, free fatty acids, or dopamine in Krebs buffer containing 1% DMSO and 25 μM Diprotin A (to inhibit DPP4) for 3–4 hours.

Supernatants were then collected and either frozen at -80°C or directly used for GLP-1 assays. A GLP-1 bioreporter cell line was generated similarly to the GPR119 bioreporter: a human GLP-1 receptor (GLP-1R) ORF was overexpressed in CRE-luciferase HEK293T cells, and activity was validated using the GLP-1 receptor agonist Exendin-4. NCI-h716 supernatants were added to the GLP-1 bioreporter cells at a final concentration of 20% in Krebs buffer, and GLP-1 activity was measured using the Bright-Glo Luciferase Reagent. The concentration of secreted GLP-1 was interpolated from Exendin-4 activation curves (GraphPad Prism10), generated on the same day using the same bioreporter cells.

### EEC-containing human gut organoids

Human gut organoids were generated using a previously described differential protocol to enrich for enteroendocrine cells (EECs)^32^. The presence of EECs was confirmed by immunofluorescence staining for chromogranin A. RNA was extracted for quantitative reverse transcription PCR (qRT-PCR) to evaluate the expression of free fatty acid receptors (FFAR1, FFAR2, FFAR3, FFAR4, and GPR119). Organoids were incubated with acyl amines in assay media, as described previously, for 2, 5, or 24 hours. Supernatants were then collected and analyzed for GLP-1 secretion using the assay described above.

### Gnotobiotic Mouse Studies

All animal studies were approved under an institutional IACUC protocol at the Massachusetts Host-Microbiome Center’s Gnotobiotic Resources. Defined-colonization experiments in gnotobiotic C57/BL6 mice were conducted in positive-pressure gnotobiotic isolators (Class Biologically Clean, Madison, WI). Mice were gavaged with 1×10^8^ CFU of *Bacteroides thetaiotamicron* (strain ATCC 29148, American Type Culture Collection, Manassas, VA) to pre-condition the gut environment, followed by oral gavage 7 days later with *Blautia wexlerae* (strain 59E11 from the GMbC) or control gavage with vehicle alone. Fecal pellets from mice were collected at 72 hours post-*Blautia* challenge and cultured anaerobically on Brucella, CAN and BBE agar (Thermo Fisher, Waltham, MA) to confirm colonization or maintenance of the GF state. Aerobically incubated plates confirmed the absence of aerotolerant contaminants. Identification of colonized bacteria was performed by colony inspection, Gram stain of fecal pellets, and amplification and Sanger sequencing of the 16S gene for at least 50 colonies per isolators. Four weeks after colonization confirmation, mice were placed on a high-fat diet TD.88137 (Inotiv; Indianapolis, IN). Mice were weighed using battery-powered balances weekly. Mice were sacrificed at 12 weeks after starting the high-fat diet. Oral glucose tolerance was performed (details below) before sacrifice. Serum GLP-1 was also measured as detailed below. Stool was collected for LC-MS, using protocols as described above.

### Oral Glucose Tolerance Test in Mice

C57BL/6J mice (aged 8-12 weeks) were fasted for 6 hours prior to the start of the oral glucose tolerance test (OGTT). Mice were divided into experimental groups (n = 8 for acyl amine treatments and n = 8 for vehicle control) or into colonization groups (see above) and baseline blood glucose levels were measured via tail vein sampling using a handheld glucometer (OneTouch Ultra). Three acyl amines were tested: oleoyl dopamine, linoleoyl dopamine and α-linolenoyl dopamine. Acyl amines gavaged at doses of 100 mg/kg and solubilized in 10% DMSO and 50% ethanol in PBS. Thirty minutes after compound gavage, baseline glucose was measured and mice were administered a glucose solution (2 g/kg body weight) via gavage. For *B. theta* and *B. wexlerae* colonized and GF mice, only glucose was administered. Blood glucose levels were then measured at 15, 30, 60, 90, and 120 minutes post-glucose administration.

Blood samples were obtained from the tail vein at each time point, and glucose levels were quantified using the glucometer. The area under the curve (AUC) for glucose was calculated using the trapezoidal method to quantify glucose intolerance (GraphPad Prism10).

### Serum GLP-1 Measurement

Mice were fasted for 4 hours prior to oral glucose stimulated GLP-1 measurements. Mice were divided into experimental groups (n = 8 for acyl amine treatments and n = 8 for vehicle control; or colonization groups as described above). For acyl amine testing, a combination of three acyl amines (oleoyl dopamine, linoleoyl dopamine and α-linolenoyl dopamine) were gavaged at a total dose of 100 mg/kg body weight, solubilized in 10% DMSO and 50% ethanol in PBS, followed by a glucose solution (2 g/kg body weight).

For *B. theta* and *B. wexlerae* colonized and GF mice, only glucose was given. Twenty and forty minutes after gavage, blood samples were obtained (100 μL per mouse) from the tail vein using capillary tubes to collect from a small nick. Diprotin A (Cayman Chemical; Ann Arbor, MI) was added for a final concentration of 25-50 μM) and the samples were immediately placed on ice. Plasma was used to stimulate GLP-1 bioreporters (see above) at a final concentration of 2% plasma in Krebs buffer to quantify active GLP-1 concentration in the blood.

### Food Intake Measurements in Mice

Mice were fasted for 4 hours prior to food intake experiments. Mice (n = 8 cages) were doubly-housed for food intake experiments. A combination of three acyl amines (oleoyl dopamine, linoleoyl dopamine, and α-linolenoyl dopamine) or vehicle controls was gavaged (total dose of 100 mg/kg body weight for acyl amines), and mice were allowed to eat ad libitum overnight (16 hours). Mice were given either acyl amines or vehicle with a 3–5-day washout period in between, for a total of three acyl amine and three vehicle control treatments. Custom-made J-shaped food hoppers with built-in spillage reservoirs were used to minimize food droppage on the cage floor. Chow was measured before and after the overnight feeding session, allowing for food intake to be measured for each cage and averaged across the two mice per cage.

## ACKNOWLEDGEMENTS

We would like to thank Barbara Kahn, Jen Nguyen, Ethan Evans, Margaret Stefater-Richards, Brian Carmine, and Anni Zhang for their thoughtful discussions. This work was supported by the NASPGHAN Foundation/Reckitt Mead Johnson Nutrition Research Young Investigator Award, the Child Health Research Career Development Award (CHRCDA) NICHD K12HD052896 and K08DK138274 to YJZ; K08DK134885 to DZ; Intramural Program of the *Eunice Kennedy Shriver* National Institute of Child Health and Human Development (ZIAHD000064) support for JAY; a capital grant from the Massachusetts Life Sciences Center and the Harvard Digestive Disease Center P30 grant DK034854 to support the Massachusetts Host-Microbiome Center (LB); the Fulbright Doctoral Dissertation Research Award grant to APS; a Springer-Nature Global Grant for Gut Health and a Charles H. Hood Foundation Child Health Research Award to MJ; and DK048106, DK122953, DK118640 and the Harvard Digestive Disease Center P30 grant DK034854 to WIL.

## Disclosures

For research unrelated to the current paper, at the time this research was performed JAY was 1) Principal Investigator for a clinical study administering colchicine or placebo to people with obesity supplied under a Clinical Trials Agreement between Hikma Pharmaceuticals, Inc and the *Eunice Kennedy Shriver* National Institute of Child Health and Human Development (NICHD); 2) Site Principal Investigator for clinical research conducted under Cooperative Research and Development Agreements between NICHD and Rhythm Pharmaceuticals (setmelanotide) and between NICHD and Soleno Therapeutics (diazoxide choline extended-release).

The contributions of the NIH authors were made as part of their official duties as NIH federal employees, are in compliance with agency policy requirements, and are considered Works of the United States Government. However, the findings and conclusions presented in this paper are those of the authors and do not necessarily reflect the views of the NIH or the U.S. Department of Health and Human Services.

## SUPPLEMENTAL TABLES

Supplemental Table 1. Differential Abundance, species level, LOC vs non-LOC eating.

Supplemental Table 2. Species with CTA gene homologs in GMBC and BIO-ML.

Supplemental Table 3. Stool Metabolomics in Colonized Mice.

Supplemental Table 4. Differential abundance testing for stool metabolites, *B. wexlerae* vs *B. wexlerae* + *B. theta* colonized mice

## DATA AVAILABILITY

The raw sequencing data generated in this study have been deposited in the NCBI BioProject database under accession number PRJNA1436573.

## Notes

### Competing Interest Statement

JAY received grant support unrelated to this article for pharmacotherapy trials for human obesity from Hikma Pharmaceuticals, Inc., Soleno Therapeutics, Inc., and Rhythm Pharmaceuticals, Inc., and reagents (anti-activin receptor antibodies) from Versanis Bio for mouse studies.

