## Supplemental Figures for "*Blautia wexlerae* Transforms Dietary Fatty Acids to Activate Enteroendocrine Signaling and Improve Metabolic Health in Mice and Humans"

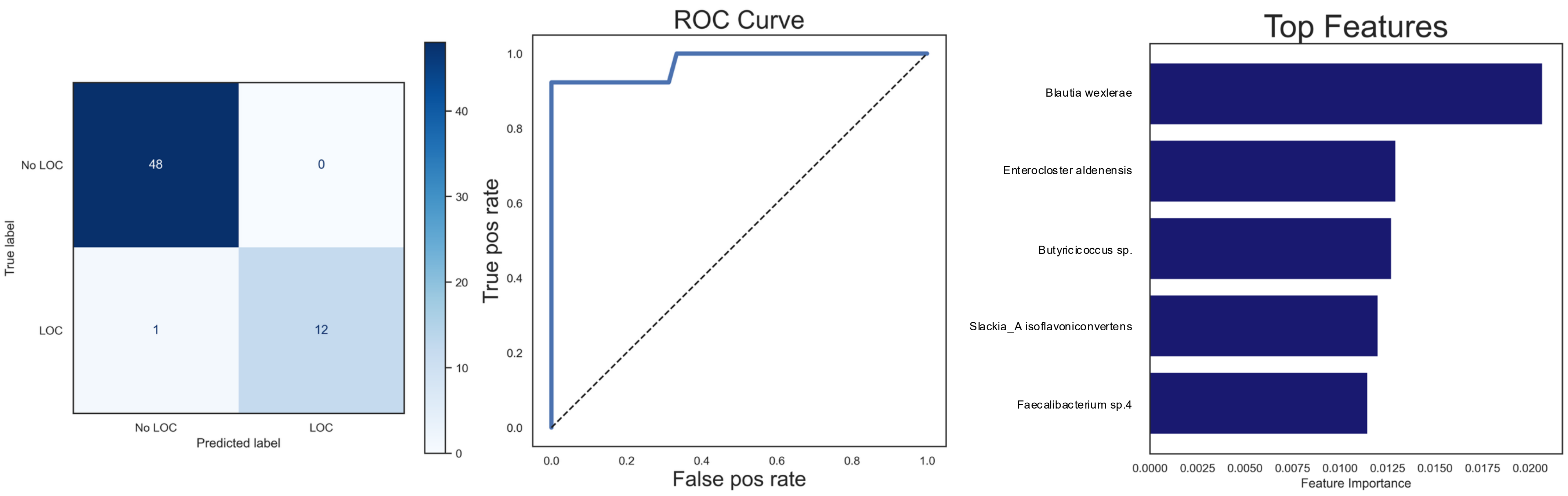


**Supplemental Figure 1.** Left: A machine learning algorithm—random forest classifier—using only microbiome data (relative abundance at the species level) accurately predicted Loss of Control in children. Due to the relative lack of participants with LOC (<10%), these participants were oversampled in both the training and test sets to about 20%. The confusion matrix demonstrates 98.3% accuracy. Center: Receiver operating characteristic curve (AUC 97.5%) of the random forest classifier showing model discrimination. **C.** The top five ranking features contributing to the random forest classifier, ordered by importance.**.**

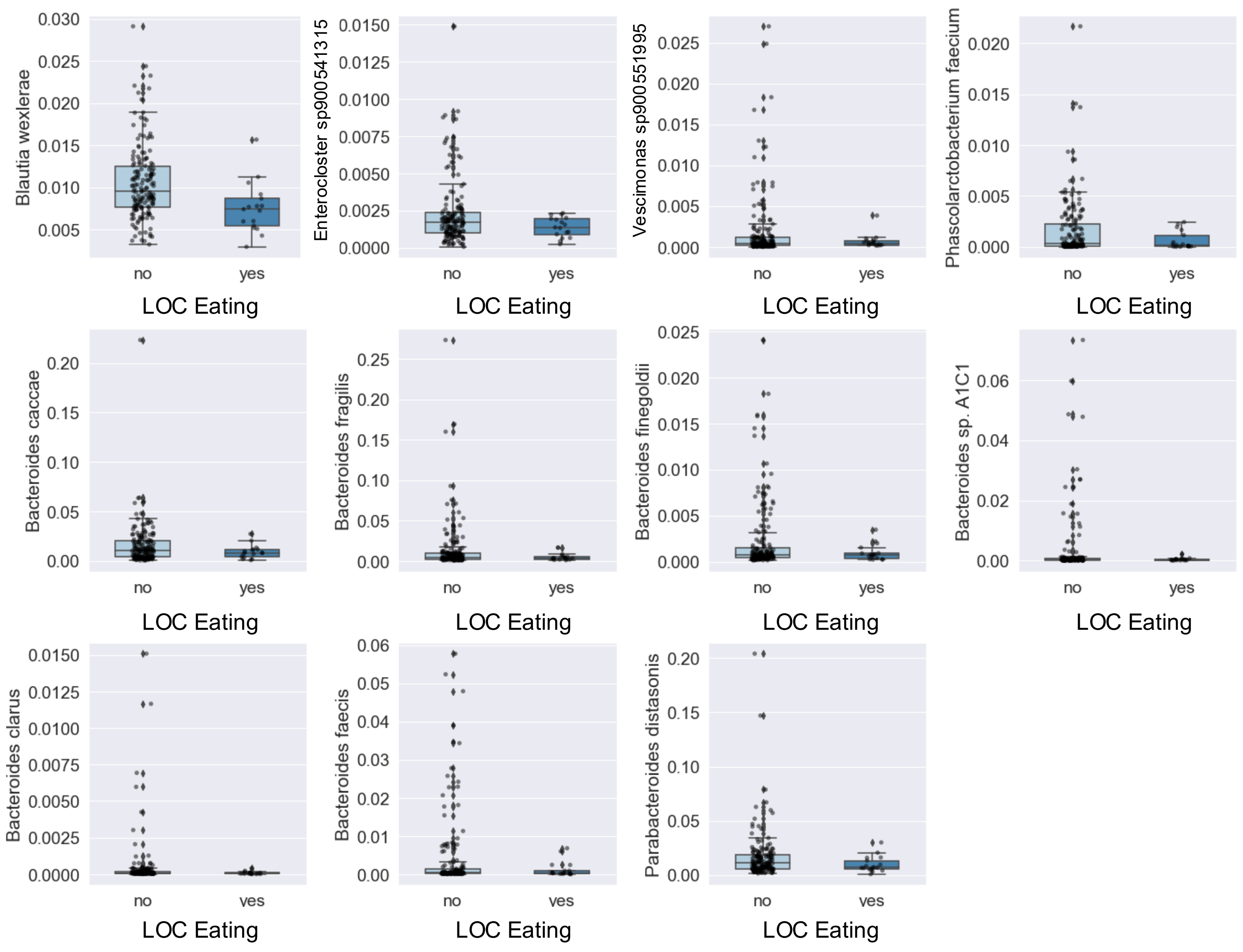


**Supplemental Figure 2.** The differentially abundant species between children with and without Loss of Control eating (unequal variance t-test, Bonferroni correction, FDR < 0.05). All differentially abundant species that made the cut-off are shown.

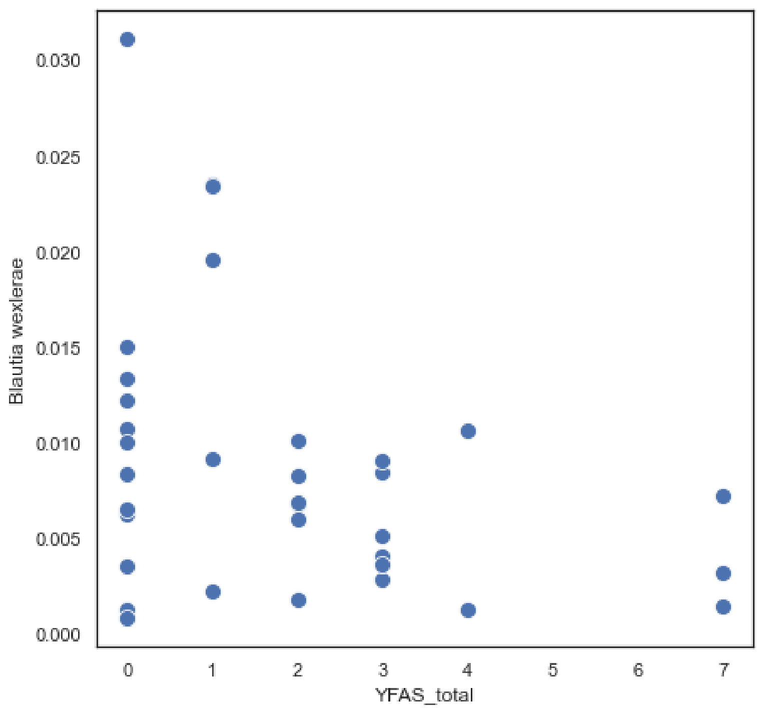


**Supplemental Figure 3.** Scatter plot showing the relative abundance of *Blautia wexlerae* in stool samples plotted against Yale Food Addiction Scale (YFAS) total scores. Each point represents one individual.

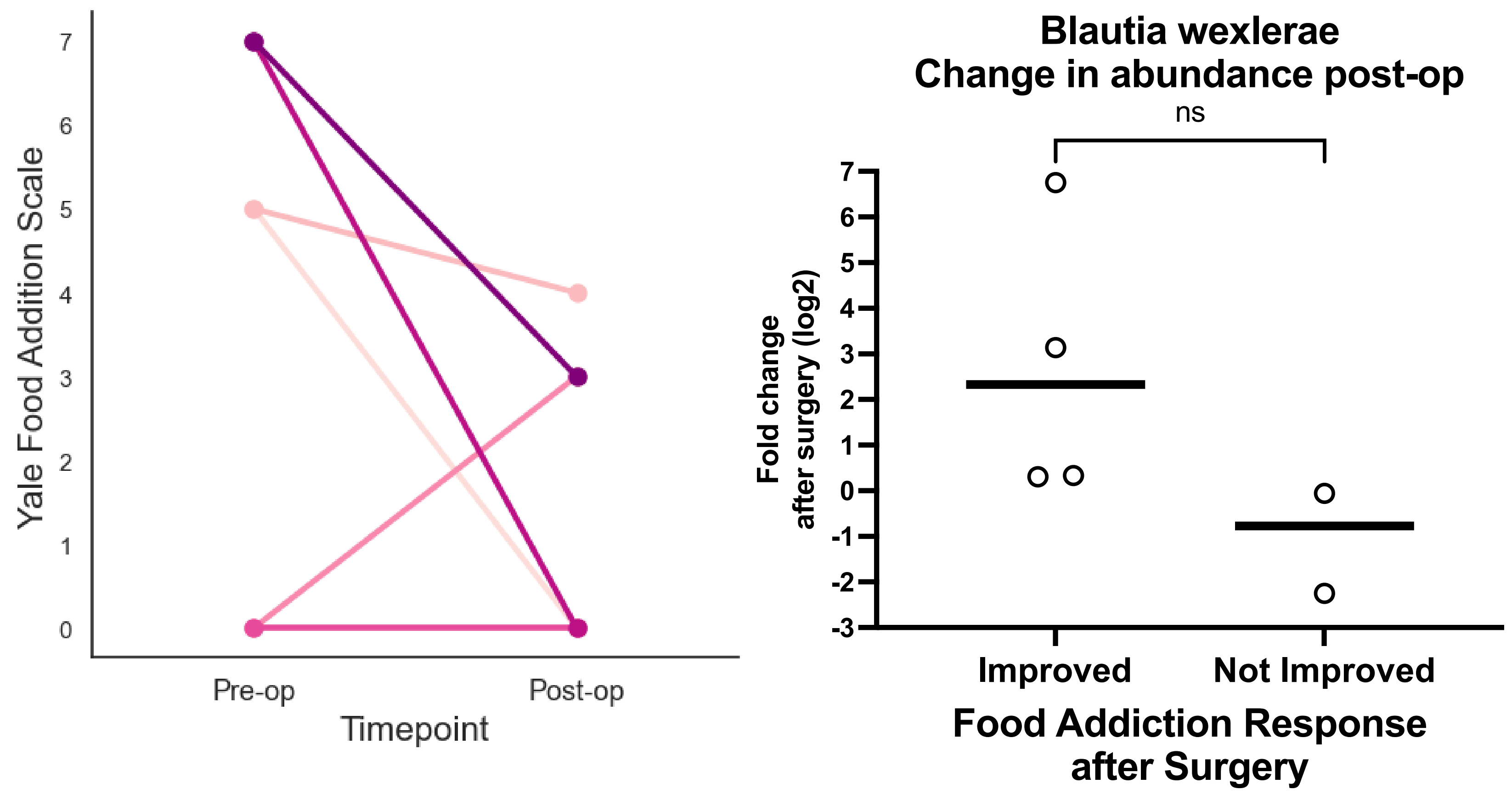


**Supplemental Figure 4.** Left. Change in the YFAS scores for patients before (3 months pre-op) and after (6-12 months post-op) bariatric surgery. Each paired dot and line represents one individual participant. Four participants (of six) experienced a decrease in YFAS scores after surgery. Right. The change in *Blautia wexlerae* relative abundance (expressed as a fold change after surgery), comparing the four participants with improved YFAS scores vs. the two patients without. On average, *Blautia wexlerae* increased in the participants with improved (less) food addiction symptoms and decreased in the participants without improvement in food addiction symptoms, though the study was underpowered and the finding was not statistically significant.

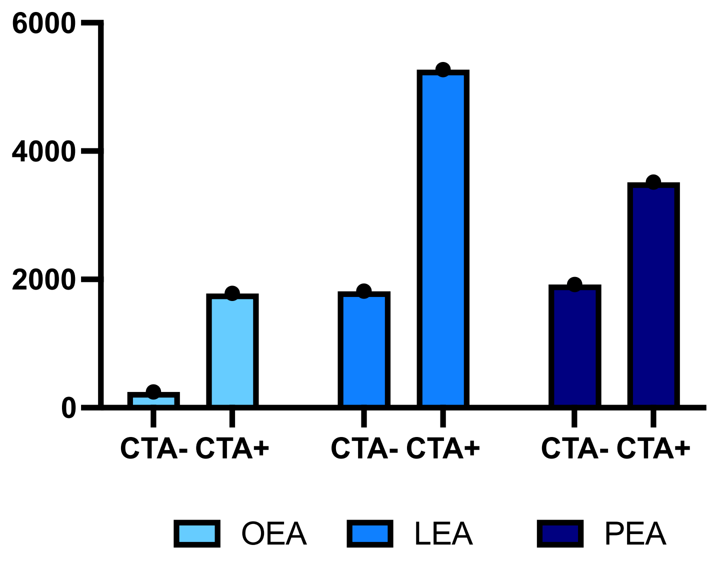


**Supplemental Figure 5.** Levels of OEA, LEA and PEA measured by LC-MS from two *Coprococcus catus* isolates, one encoding (CTA+) and one not encoding (CTA-) the operon, which we hypothesized to be at least partially responsible for acyl amine synthesis in bacteria.

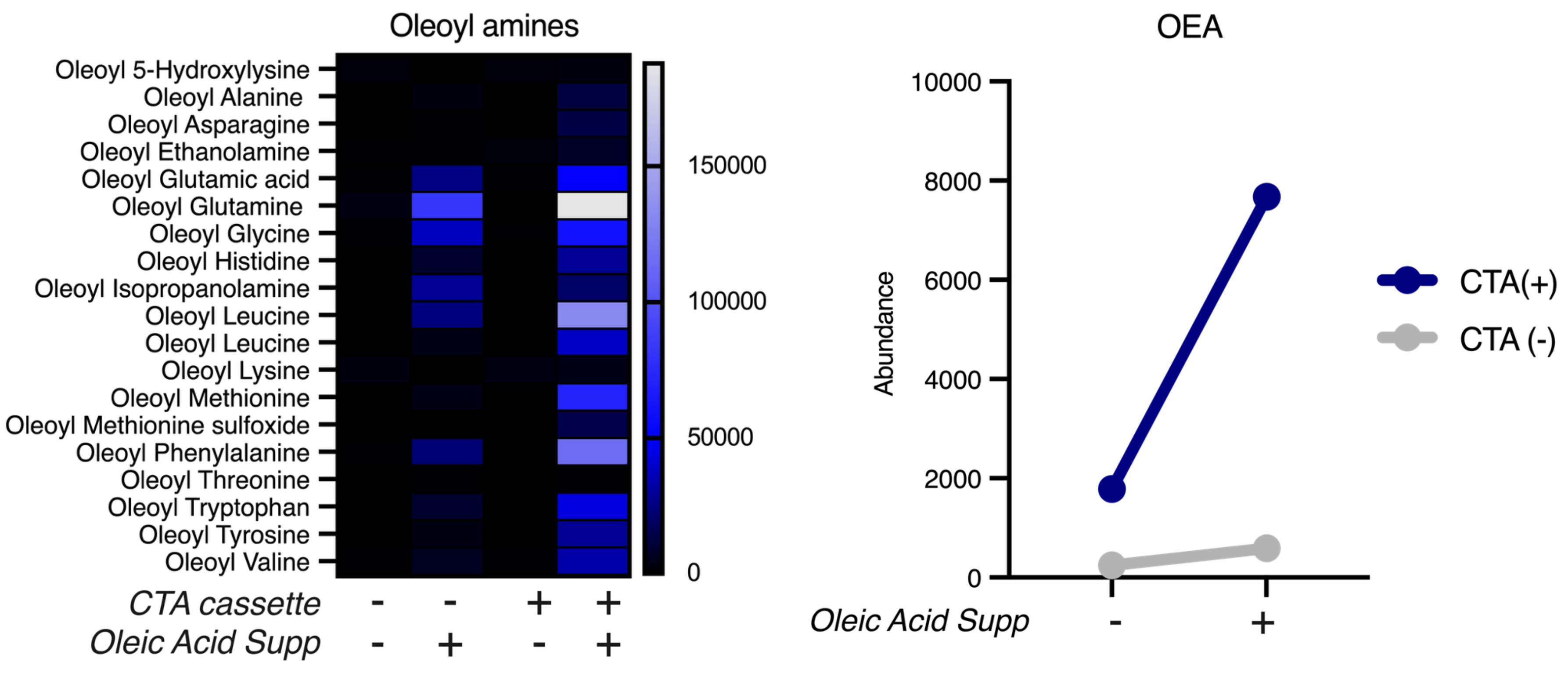


**Supplemental Figure 6. LEFT.** Two *Coprococcus catus* strains were isolated—one encoding and one not encoding the CTA operon. The *Coprococcus* strains were grown with and without supplemental fatty acids, and LC-MS was used to measure acyl amine production. Most oleoyl amines were synthesized in higher abundance by the strain encoding the CTA operon (right). **RIGHT.** The *Coprococcus catus* strain encoding the operon synthesized more OEA both before and after oleic acid supplementation.

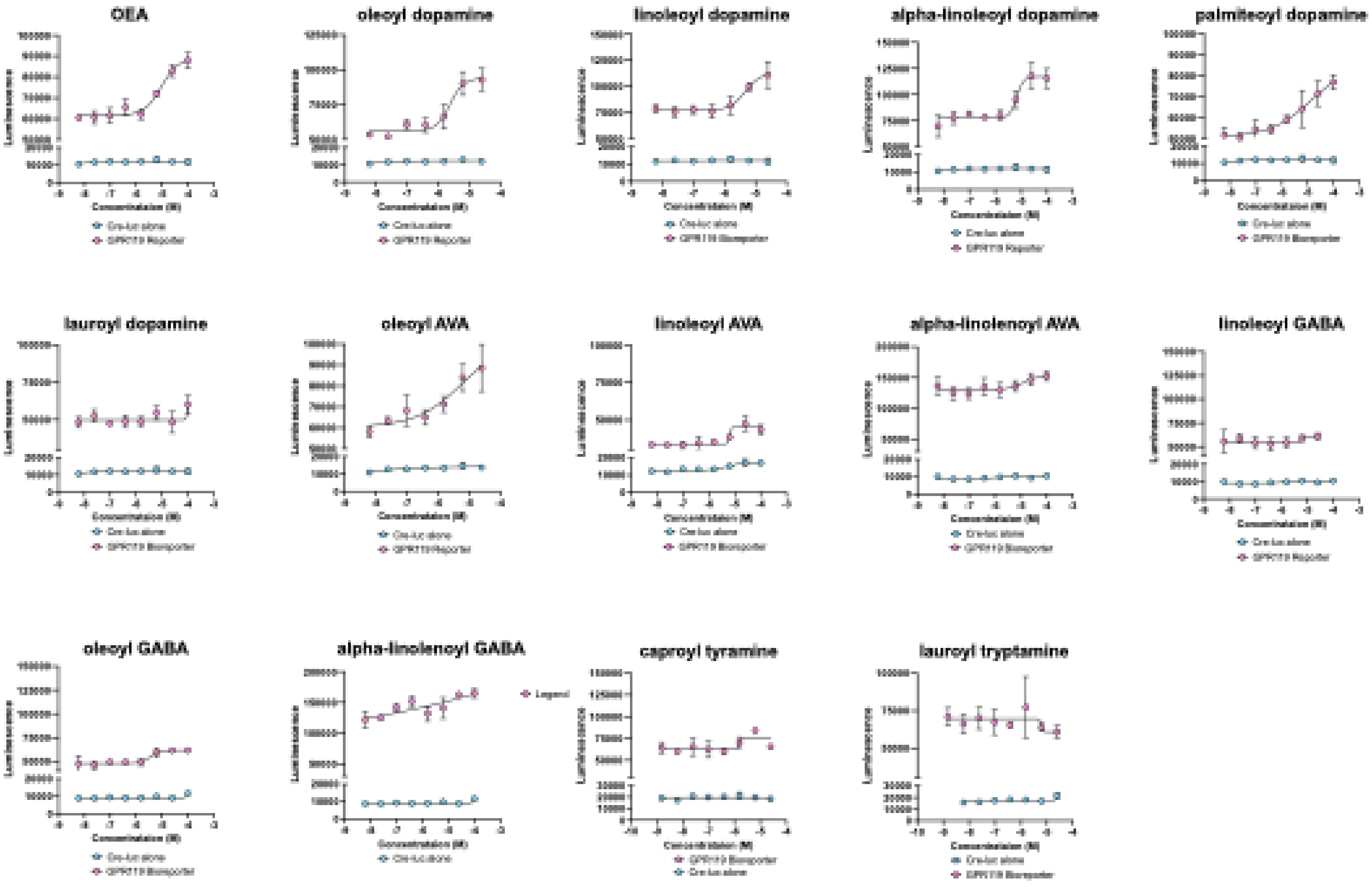


**Supplemental Figure 7.** Raw luminescence values from our GPR119 reporter cell stimulated with various acyl amines. The reporter cells are HEK293T cells expressing luciferase under the control of the cAMP-responsive element (Cre-luc) and GPR119. HEK293T cells only expressing the Cre-luc construct were used as controls to detect any non-specific activators of the cAMP. EC50s are summarized in Figure 5A).

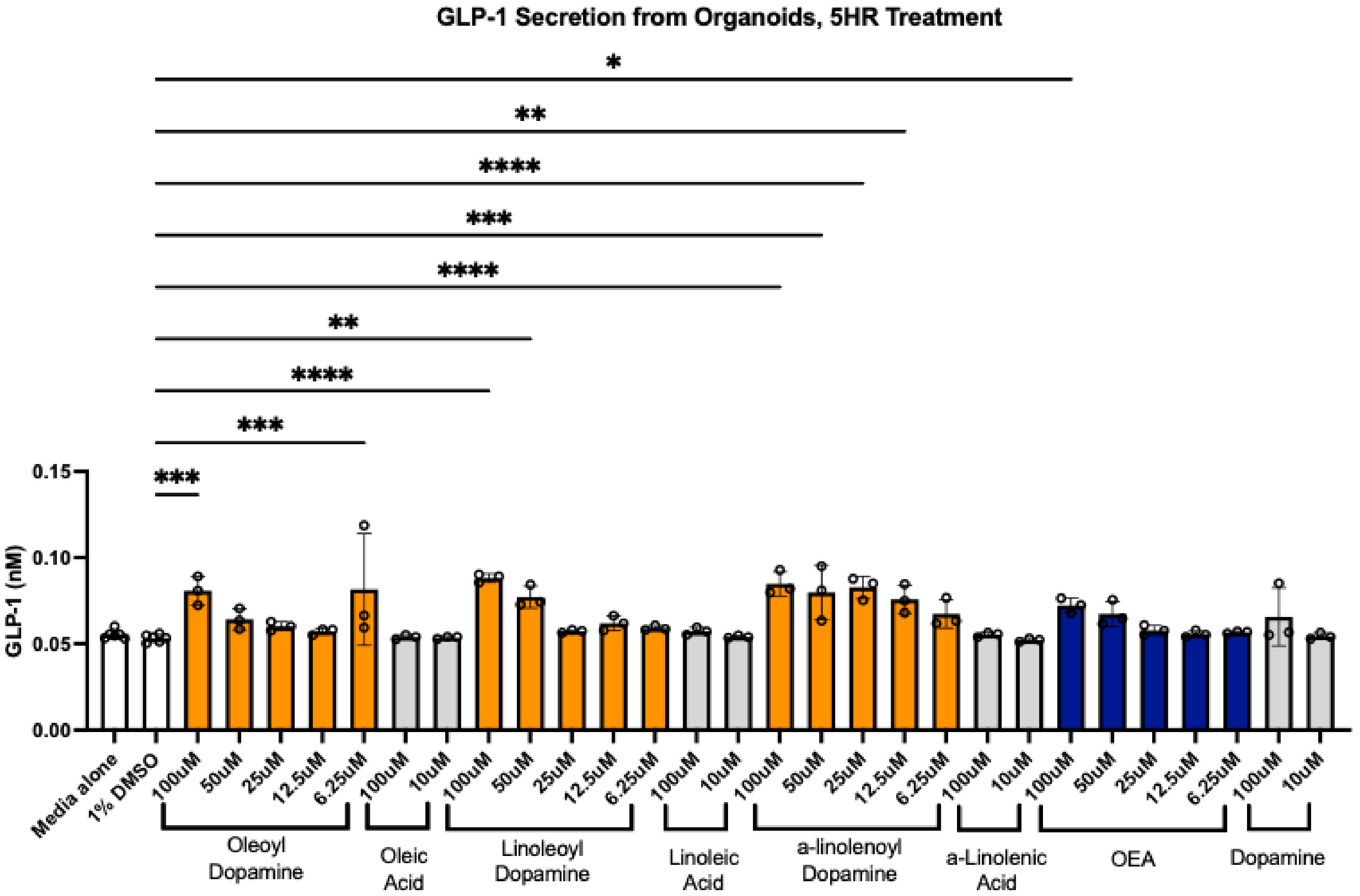


**Supplemental Figure 8.** GLP-1 secretion from EEC-containing organoids with extended dilution series, OEA, free fatty acids, dopamine and an extended (5 hr) incubation time. (*, $p < 0.05$; **, $p < 0.01$; ***, $p < 0.001$; ****, $p < 0.0001$)

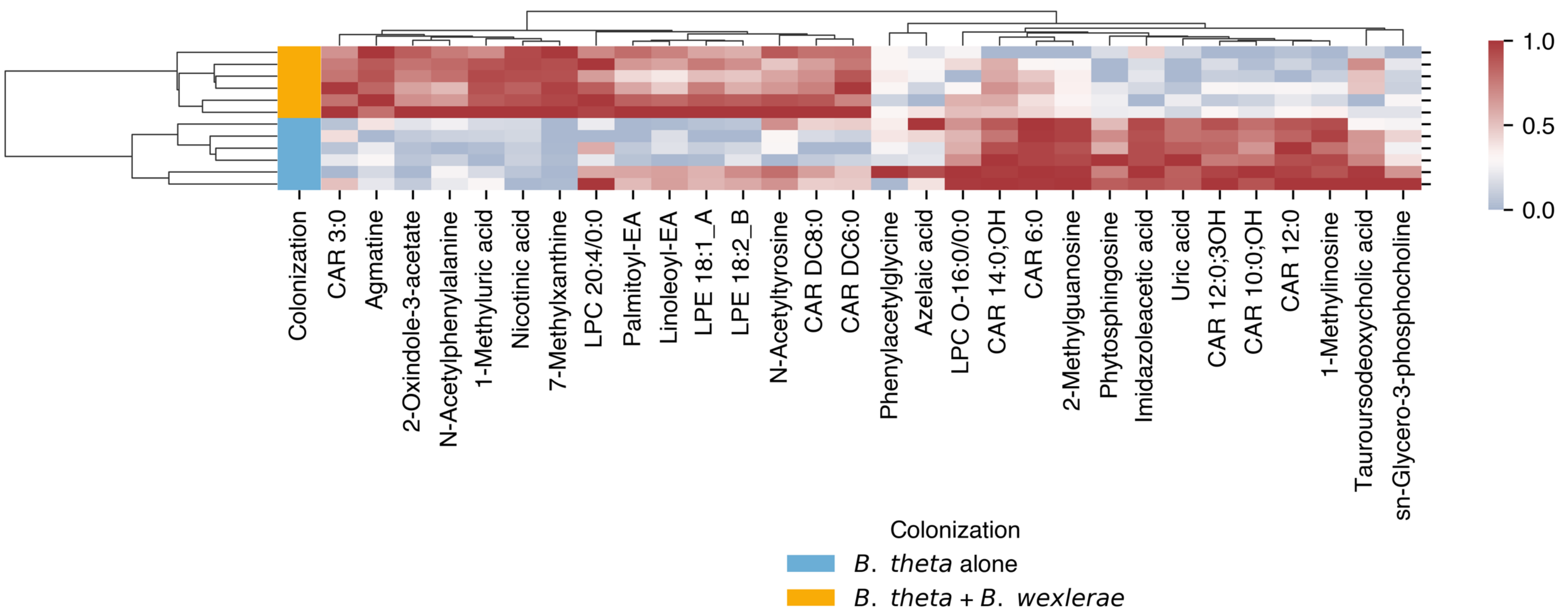


**Supplemental Figure 9.** Heat map of the differentially abundant metabolites in stool, determined by LC-MS, comparing germ-free mice colonized with *B. theta* alone vs. *B. theta* and *B. wexlerae*. All relative abundance values were normalized to the geometric mean for each metabolite. The top 30 differentially abundant metabolites are shown.

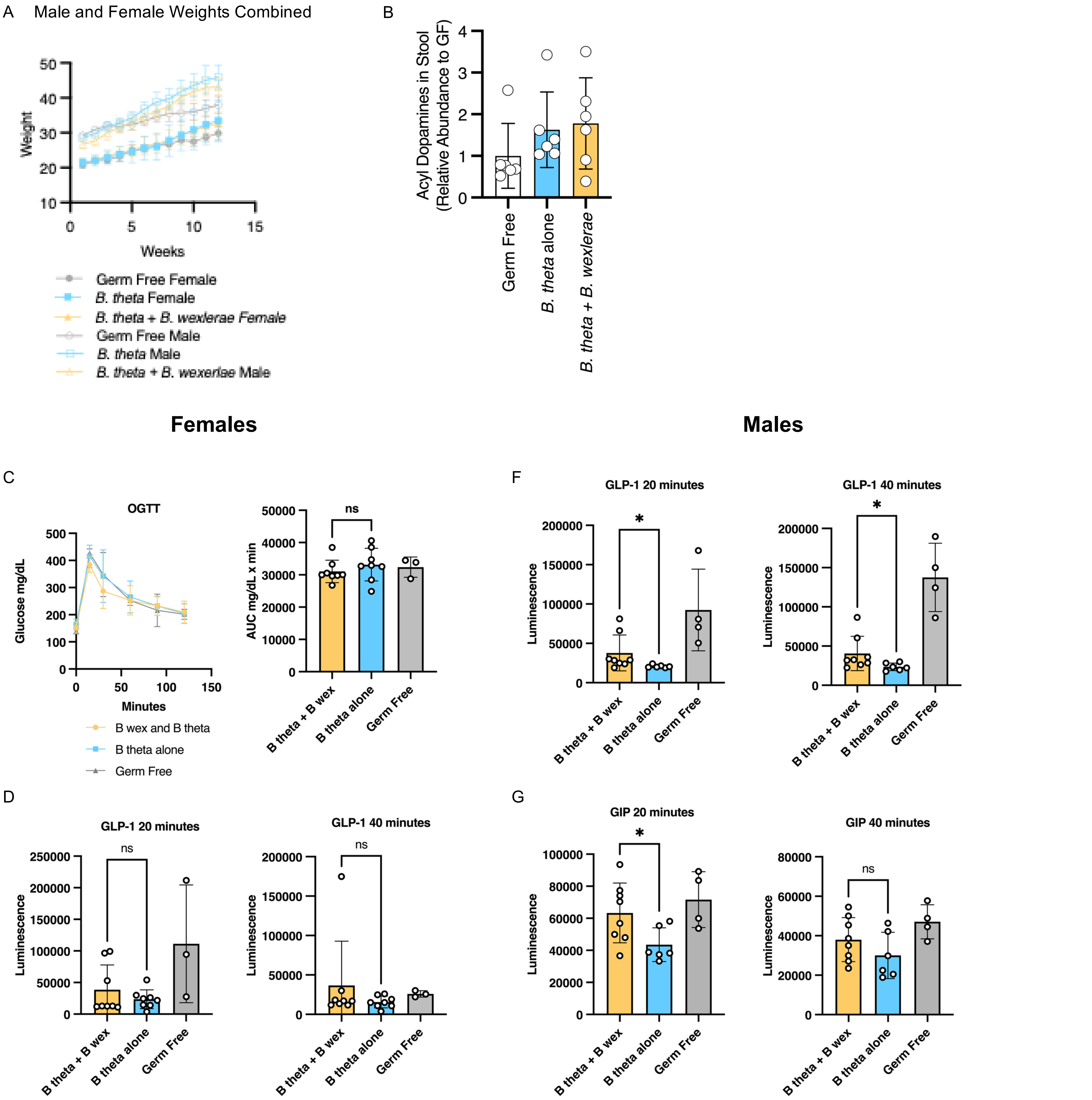


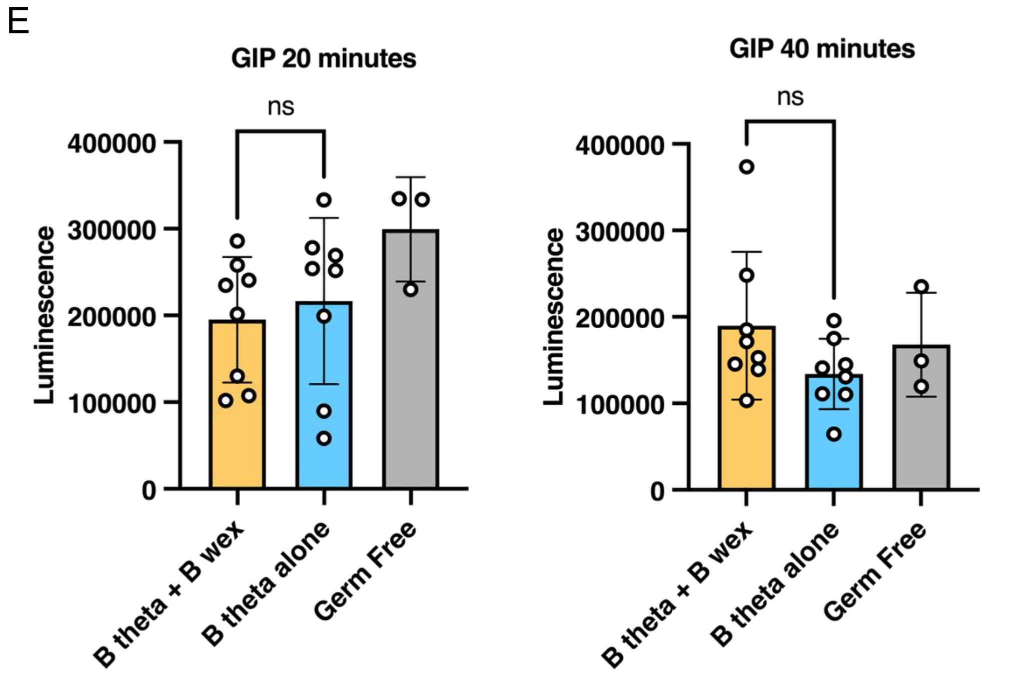


**Supplemental Figure 10.** A. Weight trends in male and female mice. Female mice were resistant to weight gain on the high fat diet, regardless of colonization status. B. Relative abundance (reference group: germ-free) of acyl dopamines, quantified from the stool by LC-MS. Mice colonized with both *B. theta* and *Blautia wexlerae* had the highest mean abundance, though this was not statistically significant. C. Oral glucose tolerance test in female mice after glucose gavage (2 g/kg). Glucose curve (left) and area under the curve (right) are shown. We observed no differences between the groups. D. GLP-1 secretion at 20 and 40 minutes after glucose gavage (2 g/kg) in females. E. GIP secretion at 20 and 40 minutes after glucose gavage (2 g/kg) in females. F. GLP-1 secretion at 20 and 40 minutes after glucose gavage (2 g/kg) in males. We observed differences between the two groups of colonized mice (Mann-Whitney U test; *, $p < 0.05$). G. GIP secretion at 20 and 40 minutes after glucose gavage (2 g/kg) in males. We observed differences between the two groups of colonized mice only at 20 minutes (Mann-Whitney U test; *, $p < 0.05$).

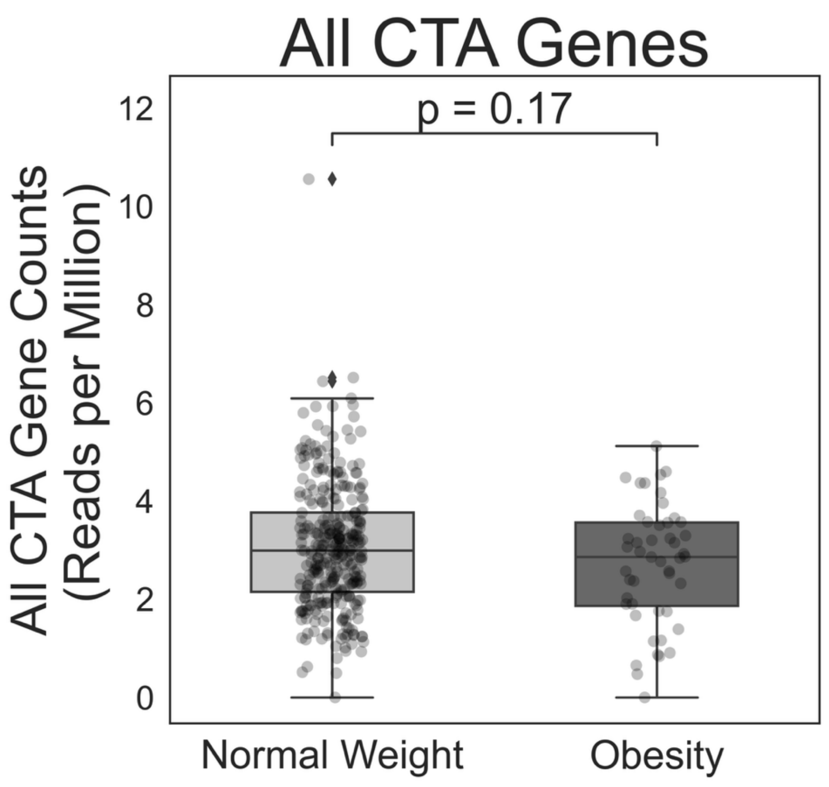


**Supplemental Figure 11.** Comparison between individuals with normal weight vs. obesity in the Global Microbiome Conservancy. Unlike in Figure 7B, here, reads were aligned to CTA genes from all *Lachnospiraceae*, rather than CTA genes from Blautia alone. There were no differences between the two weight groups.

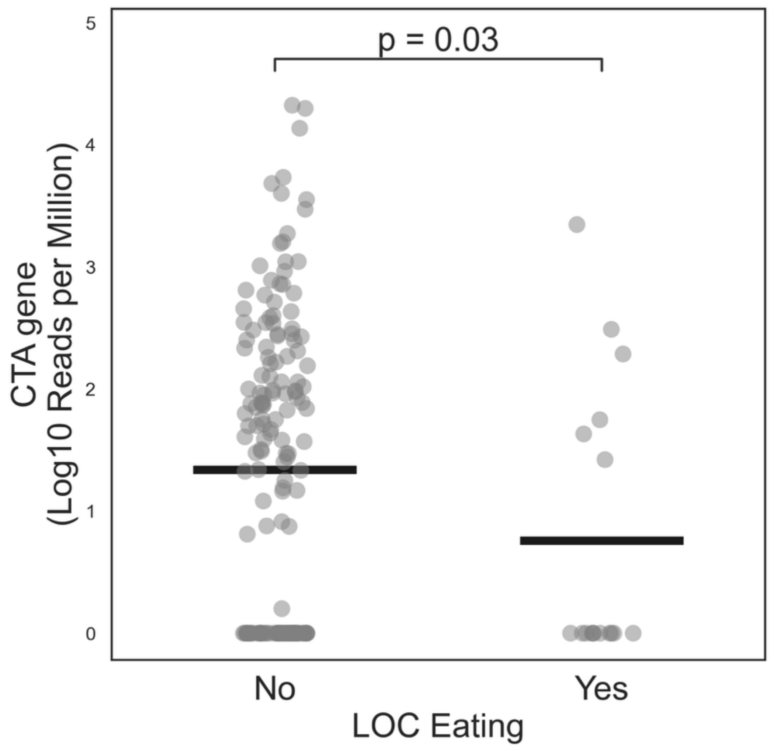


**Supplemental Figure 12.** Comparison between children with vs. without loss of control (LOC) eating. There was a significantly lower abundance of reads mapping to the *Blautia* CTA genes in participants with LOC eating (Mann-Whitney U test).
